# Ordering molecular diversity in untargeted metabolomics via molecular community networking

**DOI:** 10.1101/2024.08.02.606356

**Authors:** Elizabeth A. Coler, Alexey Melnik, Ali Lotfi, Dana Moradi, Ben Ahiadu, Paulo Wender Portal Gomes, Abubaker Patan, Pieter C. Dorrestein, Stephen Barnes, Vladimir Boginski, Alexander Semenov, Alexander A. Aksenov

## Abstract

Nature’s molecular diversity is not random but displays intricate organization stemming from biological necessity. Molecular networking connects metabolites with structural similarity, enabling molecular discoveries from mass spectrometry data using arbitrary similarity thresholds that can fracture natural metabolite families. We present molecular community networking (MCN), that optimizes connectivity for each metabolite, rescuing lost relationships and capturing otherwise “hidden” metabolite connections. Using MCN, we demonstrate the discovery of novel dipeptide-conjugated bile acids.

## Main Text

Metabolites synthesized by common enzymatic pathways share key functional motifs and structural characteristics. The constraints of biosynthesis yield not a disjointed chemical landscape, but an expansive yet organized “molecular terrain” composed of interrelated molecular families. Although we don’t know the true extent of the molecular diversity, and possibly billions of molecules may exist^1^, we can be fairly certain that underlying structural patterns of biology restrain molecules within definable “families”, preserving order amidst expansive molecular individuality. In our exploration of vast molecular complexities of biology, we can leverage this context by understanding structural relationships across metabolic landscapes.

The contextualization of molecular space to propagate structural information between related molecules has been implemented as a strategy of molecular networking, a method that operates on mass spectrometry (MS) data^2^. Mass spectrometry, in combination with chromatography (liquid, LC-MS and gas, GC-MS) is an indispensable platform for the discovery, characterization, and quantification of diverse molecules, ranging from methane to proteins^3^. The breadth of applications of MS spans elucidation of biomarker profiles that distinguish diseased and healthy states, to explore molecules produced by microbes to mediate ecological interactions and impact human health. Molecular networking harnesses the molecular context present in MS data by connecting molecules with similar spectral fragmentation patterns (tandem MS (MS/MS) for LC-MS^2,4^ and electron ionization (EI) for GC-MS^5^) into clusters. Spectral similarities translate into structural similarities, since the same substructures of different molecules tend to give rise to the same peaks in fragmentation spectra. This, in turn, enables propagating structural information from known (annotated) to unknown molecules across network connections to then glean into the “metabolomics dark matter”, i.e. molecules that are detected but do not match known structures. Currently, the molecular “dark matter” of metabolomics in a typical untargeted tandem LC-MS analysis comprises upward of 90% of observed molecular space^6^.

Leveraging molecular context has proved to be a useful approach that has resulted in discoveries of thousands of new metabolites over the past decade. As an example, molecular networking revealed a novel microbial pathway producing amino acid-conjugated bile acids through steroid core deconjugation and reconjugation^7^. Bile acids are some of the most studied metabolites due to their central role in a multitude of biological processes, from fat solubilization, to cell signaling^8^. Yet, these microbially-made molecules have eluded discovery for over a century despite intense scrutiny, and were only noticed via molecular networking.

While molecular networking has become an indispensable tool, it has inherent limitations. Conventionally, the methodology relies on arbitrarily setting global spectral similarity thresholds to link structurally similar metabolites across molecular space. Yet, the optimal connectivity is ostensibly molecular class-specific - fragmentation patterns vary greatly for different molecular families, both in variability and richness. Common threshold for all molecules may lead to fracturing molecular families and leaving large swaths of metabolites unconnected, currently, depending on the data, somewhere between 25 to 75% of detected molecular features (Figure 1e-f, S14). These features, and fractured clusters without annotated node(s) in a cluster, are not amenable to annotation propagation, thus leaving this portion of the metabolome to remain “dark”^6^.

**Figure 1.**
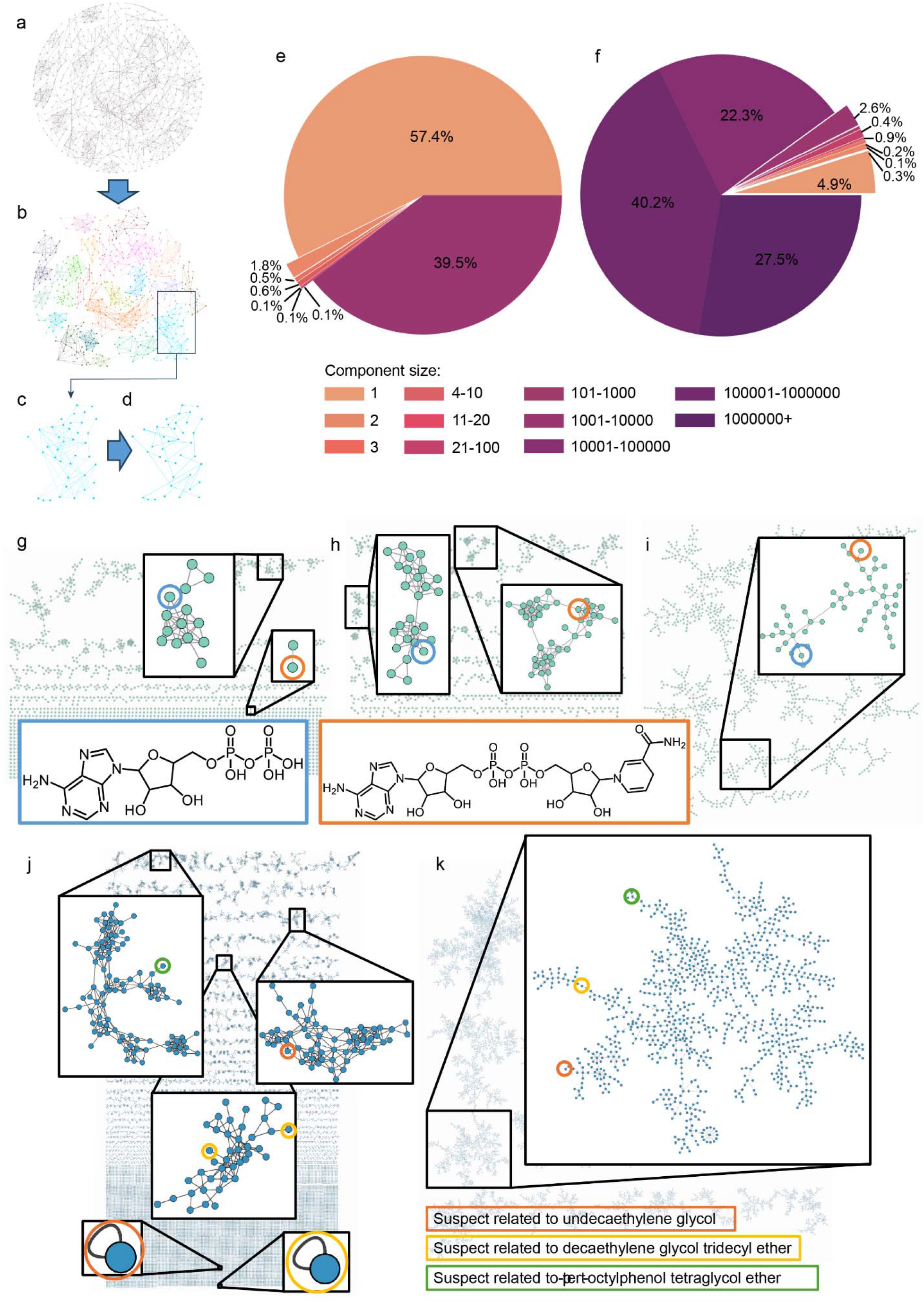
The principles and applications of molecular community networks (MCNs). Schematic description of MCN generation: **a)** An unpruned network is generated for either LC-MS/MS^2,4^ or EI GC-MS^5^ data. **b)** A clustering algorithm (Louvain method^9^) is used to determine the naturally present molecular communities within the data. **c)** Each community is pruned via the maximum spanning tree algorithm^10^. **d)** The resulting pruned community cluster is obtained by removing low-weight edges; the full connectivity is retained by keeping at least one strongest connection for each node. **e-f)** Comparison of the number of size-k connected components in conventional (e) and community molecular networks (f) for all values of k, using publicly available data from GNPS (total of ∼8.4 million metabolites). Both conventional and molecular community global networks have been generated with the MASST+ approach^29^. The number of nodes found in clusters with each size k have been counted and the percentage of the nodes within different cluster sizes is shown for the conventional network (left) and MCN (right). The coloring corresponds to the k size of the cluster, with darker colors indicating progressively larger clusters. The size-k distribution in conventional network is found to be very similar to that previously reported in^29^. The community network results in substantially higher connectivity, and exhibits approximately an order of magnitude lower number of singletons and small, uninformative clusters (2-11 nodes) compared to the conventional network. Conversely, the nodes in the community network are predominantly located within larger clusters of above 1,000 nodes, with the majority of nodes located in “superclusters” spanning thousands to hundreds of thousands of nodes. This distribution is indicative of continuity of molecular structures present in large-scale metabolomics data, i.e. molecular continuum where large numbers of metabolites are interconnected due to iterative structural variations, forming a swath of inter-related compounds rather than discrete, isolated groups. This continuity reflects the nature of metabolic processes and pathways in biological systems, where molecules often differ by small modifications or transformations, creating a vast network of structurally related compounds. In this global network of publicly available data, the additional connectivity in MCN amounts to linking over 8 million out of 8.4 million metabolites in the community network, as opposed to approximately 3.6 million connected in conventional network. **g)** A conventional molecular network generated for the reference data of 800 standards collected with different LC-MS/MS methodologies (RP, HILIC chromatography and positive, negative ionization modes), as described in Supplemental. More than one node is possible for one compound, as the same compound may be detected with different methodologies. These compounds span a variety of molecular families. A typical cosine connectivity threshold of 0.7 is used. The large portion of the molecular features remains unconnected (single nodes at the bottom of the network); positive and negative ions tend to cluster separately. **h)** The conventional network for the same reference data as in i), with the cosine threshold of 0.5. Even with such a permissive threshold, a significant portion of molecular features remain unconnected. **i)** MCN for the same data as in g) and h). Molecular families are assembled into distinct communities comprised of features detected across all methods. For the networks shown in panels g), h) and i), two members of the family of electron carrier metabolites central in the cell’s RedOx reactions are highlighted. The two shown metabolites are structurally similar, but fall into different clusters in conventional networks (g),h)); other related molecules span several other clusters. The highlighted molecules, along with all of the other various structurally related molecules all are within the same community cluster in MCN i). The close up view of these clusters is shown in Figure S9-11. **j)** A conventional molecular network generated from the LC-MS/MS data of the indoor metabolome of a house, the HOMEChem study^15^. The insets show clusters where different variants of polyethylene glycol (PEG) molecules can be found (the PEG analogues have been identified via the use of the propagated nearest neighbor, “suspect” library^16^). Usage patterns of PEG-containing personal care items leads to sizable and persistent indoor presence^15^. In a conventional network, various versions of PEGs span several clusters and many remain entirely unconnected (only a few clusters and single nodes containing PEG “suspects” are shown for visual clarity), which obscures the diversity of this molecular family in the indoor metabolome. **k)** MCN of the data shown in j). In MCN, PEG variants form a molecular community that spans molecules fractured to different clusters in j). Gathering the molecular family in a single cluster helps to better appreciate the variety of PEGs in indoor metabolome.

To resolve this challenge, we present a new rendition of the molecular networking methodology. This approach utilizes full connectivity metabolite networks that are parsed using network science tools to determine naturally present “molecular communities”, via optimization of connectivity patterns for each metabolite class. We then construct networks that preserve intra-community connectivity information. We call this approach “Molecular Community Networking” (MCN).

In conventional molecular networking, the network is pruned to keep only the connections that surpass an arbitrary cosine similarity cutoff of 0.7 or similar. In the MCN approach, we consider the *unpruned* network, with all the available information contained from all edges within this network. Then, we deploy the clustering algorithm, Louvain method^9^, on the unpruned network to detect molecular communities, i.e. groups of metabolites that exhibit high mutual connectivity (Figure 1a,b, S1). The number of communities identified by the Louvain method is not fixed or predefined; instead, the algorithm finds communities that arise “naturally” from the input data, without any initial restrictions on the number or sizes of those communities.

For networks of complex metabolomes, the natural density of links is very high, as many similar molecules are present, leading to “hairballs” - convoluted and difficult to interpret clusters. Conventionally, the “hairballs” are pruned by artificially limiting the number of connections to ∼10 for each node and limiting cluster size to ∼100 nodes. Such an arbitrary pruning without considering local patterns may remove important connectivity information. In MCN, clusters are pruned to enable network interpretation, but preserve connectivity (avoid disconnecting any nodes), while retaining only the most meaningful information. We treat each community as a separate network and identify the *maximum weight spanning tree*^*10*^ for each of these clusters. A spanning tree is a subnetwork that does not contain cycles and connects all the given nodes with a minimum possible number of links (detailed methodology is described in SI). The weight of each link represents the similarity score; thus, the maximum weight spanning tree for a network cluster would represent each cluster with only the most important links necessary to maintain full connectivity (Figure 1c,d, S1). The MCN is thus a partition of the entire original unpruned network into innate communities with a continuum of molecular space within, where the connections represent the most similar pairs of metabolites across the entirety of detected metabolome. Consequently, MCN connects nodes that otherwise may not have been connected by conventional molecular networks. As an example, on a “global” network of all public data across the GNPS/MassIVE repository^2^, increased connectivity captured by MCN leads to linking of ∼95% of all nodes (over ∼8 million out of approximately 8.4 millions of metabolites). Compared to the conventional network, this translates into additional ∼3.5 millions of nodes connected to an annotated node and thus becoming amenable to annotation propagation. This is, in large part, due to the reduction from 57.4% of unconnected nodes (“singletons”) in the conventional network to 4.9% in MCN (Figure 2e-f). We show that these connections are meaningful and link structurally-related molecules (Figure S17). It is important to understand that an MCN connection does not guarantee that metabolites are closely structurally related. Rather, it indicates that no other more similar MS/MS spectra can be found in the data. As with conventional molecular networks, a researcher’s best judgment is essential when interpreting the network.

**Figure 2.**
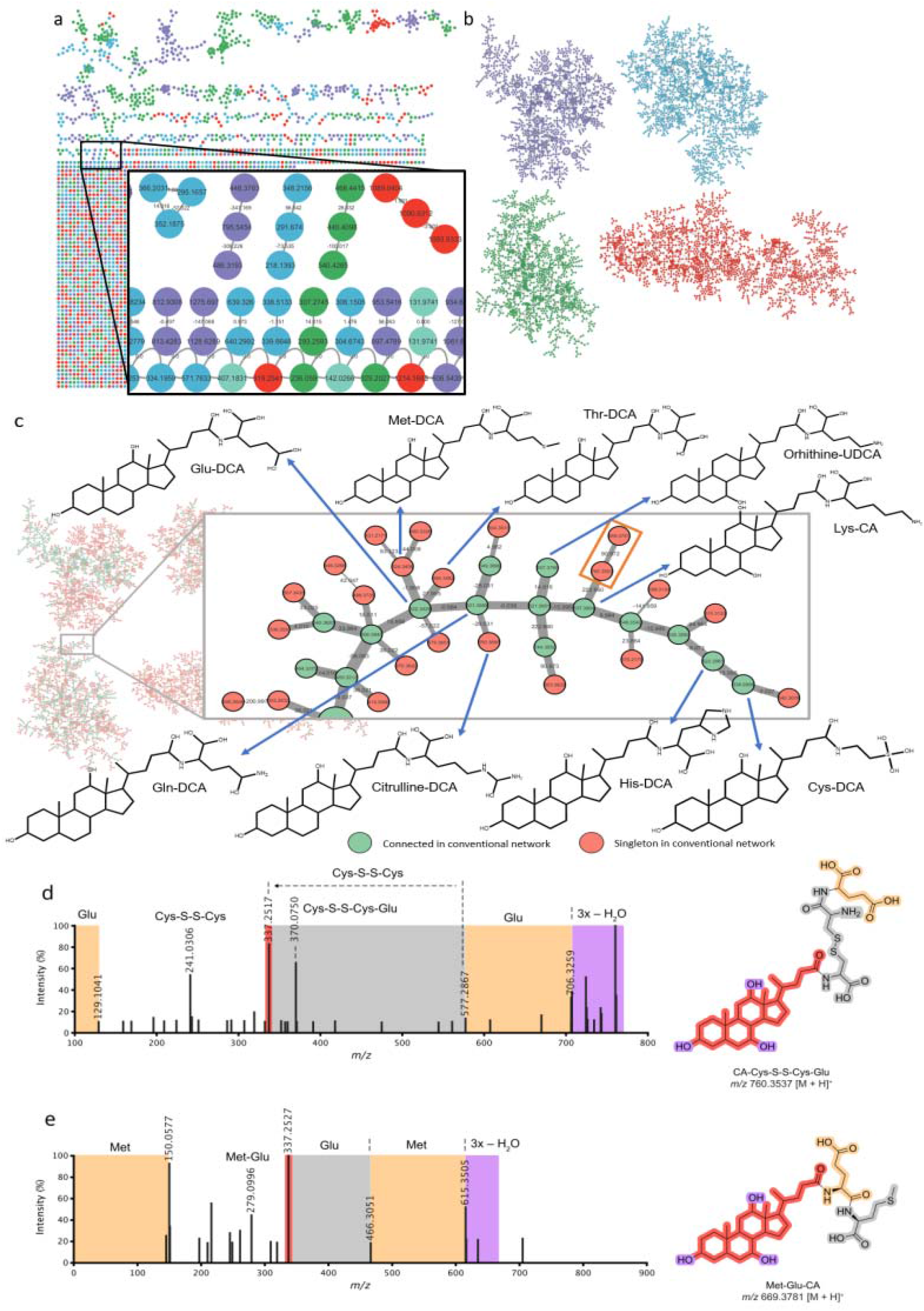
Discovery with molecular community networks. **a)** A conventional molecular network generated from LC-MS/MS data for the analysis of human microbial cultures in a study of reverse metabolomics as a discovery strategy^17^. The total of 202 isolates from skin, gut, and vaginal microbiomes were tested for bile acid conjugation capabilities. The resulting molecular distribution comprised an array of structurally dissimilar molecules, leading to a very low connectivity in the conventional network, as shown on the inset. As much as 80% of all MS/MS comparisons of bile acids have cosine similarities below 0.7 threshold and are not amenable to structure propagation in conventional molecular networks^18^. **b)** MCN of the data shown in a). The coloring in both panels a) and b) corresponds to the identified molecular communities. **c)** An up close view of a portion of a molecular community that contains bile acid molecules where the cholate core is conjugated to various amino acids, the new class of microbially-made metabolites^7^. The edge thickness is proportional to the cosine similarity score; the color indicates whether the node has any connection in the conventional network (green, connected; red, not connected; the connection may be in any cluster across the network). Proposed structures* from library matching and annotation propagation are listed for nodes across the cluster. **d)** The proposed structure* for the unannotated node with *m/z* 760.3537 highlighted by the orange box on panel c). The structure features an unexpected disulfide bond between cysteines instead of the peptide bond. This molecule is absent in public data outside of the culturing experiment**, suggesting that it is likely of artifactual, rather than biological origin. The formation of this structure provides a plausible explanation for the scarcity of free cysteine amidates across biological data, as air oxidation would have been sufficient to generate the dimers. **e)** The proposed structure* for the unannotated node with m/z 699.3781 highlighted by the orange box on panel c). Based on the MS/MS spectrum, the structure is proposed as Met-Glu-CA. An example of a dipeptide-conjugated bile acid, a glycine-taurine conjugate, has been previously reported in rabbits^19^. However, the proposed structure, featuring a different dipeptide conjugation to a steroid core, has not been described before. This structure is present in biological data** (Figure S15, S16), can be found in infant fecal samples and is associated with *Bifidobacterium breve* and other unclassified *Bifidobacterium* species. ^*^ The structural confirmation at level 2 has been carried out as described in the Supplemental^20^. ^**^ The detailed description of structures searched in public data is given in Supplemental.

We show how MCN allows assembling the molecular space for both LC-MS/MS and EI GC-MS data to reflect the continuum of structures. We explored MCNs for the reference spectra (Figure 1g-i, S2-11) to establish the validity of the connection in generated networks, and experimental data to showcase extraction of insight from the data. In one example, we show how MCN rescues lost connectivity between related molecules that conventional networking fractures into separate clusters (Figure 1j,k). In another example, we show how MCN links sodiated ion variants to the corresponding protonated precursors (Figure S12,13). Because metal adduction alters fragmentation, these features are disconnected in conventional networks and post-hoc analysis is needed to identify different versions of the same molecule^11^. However, spectra of sodiated metabolites are still more similar to the corresponding parent compounds than any other molecules, which is captured by MCN.

Linking previously stranded nodes allows propagating structural information. We demonstrate this utility of MCN by showing discovery of new bile acid structures, with dipeptite conjugation to the steroid core produced by human microbiome (Figure 2, S15-16), revealing the metabolic capacity of the human microbiota to synthesize these compounds. One of these molecules appears to be present in the stool of infants and is associated with *Bifidobacterium breve* and other unclassified *Bifidobacterium* species (see Supplemental for details). Such molecules have lower similarity connections that could not be captured by conventional networks.

Finally, we note that for all of the examined data, both LC-MS/MS and EI GC-MS (Table S1), the MCNs consistently exhibit high modularity (the metric of the ratio between “intra-cluster” vs. “inter-cluster” links; high modularity is indicative of stronger community structure). This is suggestive of the natural tendency of molecules to group into communities in metabolomics networks. We further note that molecular connectivity patterns in MCNs exhibit so-called “small-world” structure^12^ with the power-law distributions of degree of connectivity - a distinctive shape close to a straight line in the logarithmic scale (see Supplemental Figure S18), in a notable similarity to that of online social networks^13^ and other connected systems. Such connectivity described by power law distributions can be found in diverse domains, including languages, neuronal firing patterns, organization of firms, protein-protein interaction networks, etc. These systems are characterized by self-organization into hierarchical structures with functional modularity (organization into distinct, semi-independent units or modules that perform specific functions). The similarity of molecular networks to these networks may further enable deeper biological insight by transposing known properties and patterns in various systems. The topology of these networks may be informative in itself, and reveal informative patterns of molecular space in general, such as identifying the most “central” nodes, groups of nodes, or paths in these networks^14^. We anticipate this approach to empower molecular discovery, in areas such as natural products research, including reanalysis of existing data to explore molecules previously unconnected in conventional networks.

## Data availability

All data used for testing and validating molecular community networking are deposited in GNPS/MassIVE^2^. All data and links to the GNPS jobs underlying figures present in the Main Text and Supplementary Note are included as Supplementary Data.

## Code availability

We have deposited the code at github https://github.com/Alexander0/molecular_communities. The molecular community networking notebook is written in Python. It is open source and released under an LGPL-3 license.

## Acknowledgements

AAA, AL were supported by the USDA NIFA GRANT13665683. PCD was supported by R01DK136117.

## Author information

### Contributions

AAA proposed the concept of molecular community networking. AS, VB devised mathematical underpinning of the methodology. AS performed the code engineering to enable molecular community networking. AAA, AL, DM, EAC curated data used in analyses. AAA, AL, DM, EAC, AVM generated data for the manuscript. WG, AM, SB established structures for the novel bile acids. PCD, WG, AP, BA conducted synthesis of bile acids to verify the proposed structures. AVM performed the reference data collection for Figure 1e-f. AAA, EAC generated unpruned networks for all data. AS, VB tested the molecular community networking code. AAA provided supervision and funding for the project. AAA, VB, AS wrote and edited the manuscript. All authors reviewed and approved the manuscript.

## Ethics declarations

### Competing interests

AAA, and AVM are founders of Arome Science, Inc. and BileOmix, Inc. PCD is an advisor and holds equity in Cybele, BileOmix and Sirenas and a Scientific co-founder, advisor and holds equity to Ometa, Enveda, and Arome with prior approval by UC-San Diego. PCD also consulted for DSM animal health in 2023.

Supplementary Note

Supplementary Table S1

Supplementary Table S2

Underlying data for the Figures

Underlying data for Figures in the Supplementary Note.

