## Supplemental Note for "Ordering molecular diversity in untargeted metabolomics via molecular community networking"

Mapping Complex Metabolomes With Molecular Community Networking

#### Overview

The making of MCN is conducted in three steps (Figure 1a-d, Figure S1):

1. Generation of an unpruned network with all possible links between all nodes (Molecular Networking).
2. Detection of molecular communities in the data (Louvain Community Detection Algorithm).
3. Pruning of sub-network of each molecular community to generate MCN (Maximum Weight Spanning Tree).

These steps are described in detail next.

#### Molecular Community Networks

As the molecular space, especially for biological systems, is not random, but contains continuity stemming from conserved molecular building blocks/substructures, any given metabolite is likely to reside amongst a neighborhood of structurally related molecules. The molecular community network is designed to capture this molecular space and generate the pairwise connections for metabolites with most similar tandem MS (MS/MS) or electron ionization (EI) spectra. The network, therefore, does not contain “singletons” (unconnected nodes), and at least one connection is present for each node. In molecular community networks, connected metabolites represent the highest spectral similarity associations discoverable within the dataset. However, for sparsely populated chemical spaces, the "nearest neighbor" for a given molecule may demonstrate only tangential structural relatedness and thus a lower cosine score. Such long-range connections likely arise when no closer biosynthetic relatives are detected, rather than indicating a definitive biochemical relationship. Therefore, while linking all possible molecules aids visualization of chemical interconnectivity, discretion is essential for annotation propagation across links with cosine scores substantially below intra-family levels, especially below cosine similarity score of ~0.6 (Figure S18). Structural inferences are generally reliable for connections that align with known compound classes. However, outliers that differ significantly from expected patterns should be interpreted cautiously. It is recommended to omit considering any connections if they fall below cosine of ~0.4 (see Figure S17).

##### Unpruned Networks Generation

We have conducted a systemic evaluation of effects of settings on production of MCN. Correspondingly, we outline the following general steps to generate the molecular community networks as follows.

- Launch a networking job in the GNPS ecosystem (<https://gnps.ucsd.edu/>)^1^ using appropriate workflow for:

Classic LC-MS:

<https://gnps.ucsd.edu/ProteoSAFe/index.jsp?params=%7B%22workflow%22:%22METABOLOMICS-SNETS-V2%22,%22library_on_server%22:%22d.speclibs;%22%7D>

Feature-Based LC-MS:

<https://gnps.ucsd.edu/ProteoSAFe/index.jsp?params=%7B%22workflow%22:%22FEATURE-BASED-MOLECULAR-NETWORKING%22,%22library_on_server%22:%22d.speclibs;%22%7D>

GC-MS:

<https://gnps.ucsd.edu/ProteoSAFe/index.jsp?params=%7B%22workflow%22:%22MOLECULAR-LIBRARYSEARCH-GC%22%7D>

The recommended network settings are listed below (under “Advanced Network Options”). However, researchers should use their discretion to select the most appropriate settings for each specific dataset, as the optimal parameters may vary depending on the nature and complexity of the data being analyzed.

- Large datasets (over ~10,000 features):

Min Pairs Cos: 0.3**

Network TopK: 100

Minimum Matched Fragment Ions: 5 or above*

Maximum Connected Component Size (Beta): 0 (disable)

Maximum shift between precursors: Use Default

- Medium datasets (~3,000-10,000 features):

Min Pairs Cos: 0.1**

Network TopK: 1000

Minimum Matched Fragment Ions: 5 or above*

Maximum Connected Component Size (Beta): 0 (disable)

Maximum shift between precursors: Use Default

- Small datasets (under ~3,000):

Min Pairs Cos: 0.01**

Network TopK: 10000

Minimum Matched Fragment Ions: 3 or above*

Maximum Connected Component Size (Beta): 0 (disable)

Maximum shift between precursors: Use Default

All other networking inputs (e.g. libraries, inclusion of metadata etc.) should be the same as for generation of conventional networks. The detailed instructions on launching GNPS job(s) are given here: <https://ccms-ucsd.github.io/GNPSDocumentation/quickstart/>

*NOTE: Setting the “Minimum Matched Fragment Ions” too low could result in spurious connections.

**NOTE: It is not possible to set “Min Pairs Cos” at 0 value; therefore, an alternative, small value is used to generate the unpruned network. The suggested values are listed above. For sparse data with heterogeneous molecular composition, it is possible that even at such low similarity values some features will be unconnected (singletons). These features are not included in the downstream MCN generation.

##### Community Network Generation

The molecular community network is generated from the unpruned network. Once the GNPS job for the unpruned network is finished, the following sequence of steps is needed to generate the community network:

- From GNPS job page, download GraphML file using the corresponding link:
  - “Download GraphML for Cytoscape” (classical LC-MS network)
  - “Download Cytoscape Data” (FBMN LC-MS network, GC-MS network)
- Using downloaded graphML file, launch the notebook downloaded from [https://github.com/Alexander0/molecular_communities](https://github.com/Alexander0/molecular_communitiescode)
- The notebook requires the 'networkx' library v. 3.2.1, which can be installed using pip. To run the notebook, modify three variables: INPUT_FILE_PATH, OUTPUT_PATH, and OUTPUT_NAME. They should point to the input GraphML file, the folder that will store the resulting networks, and the name of the file used for output generation, respectively.
- The molecular community networking files will be generated in the folder OUTPUT_PATH. Three files, the unpruned network with all possible connections retained, network pruned by the Spanning Tree and Spanner/Spanning Tree will be placed in the same folder as the input network file.
- Download the molecular community networking graphML file: “OUTPUT_NAME_spanning_tree.graphml”. This file can now be used for generating molecular networks, using any suitable software such as Cytoscape (https://cytoscape.org/). The tutorial could be found here: <https://ccms-ucsd.github.io/GNPSDocumentation/cytoscape/>. The visualization can be tailored to the specific analyses as with conventional networks.

##### Louvain Community Detection Algorithm

The Louvain Community Detection Algorithm^2^ provides a straightforward and efficient approach for extracting the community structure within a network^3,4^. This method centers around modularity optimization. The Louvain algorithm is a heuristic method: it produces a high-quality albeit not necessarily the exact optimal solution (in terms of maximum modularity), and it provides substantial practical advantages compared to exact optimization approaches for community detection that lead to computationally challenging (NP-hard) network optimization problems^5–9^. Such exact optimization problems typically cannot be solved within a reasonable time for large networks. Conversely, the Louvain algorithm performs very well in terms of both the quality of the solution and the computational performance/scalability, as it can successfully identify communities via modularity optimization in very large networks (e.g., networks with ~100M nodes^10^). The high scalability of the Louvain algorithm makes it especially suitable for Molecular Community Networking in very large molecular networks, potentially with millions or even billions of molecules.

Network modularity is a measure that quantifies the degree to which a network can be naturally divided into distinct, internally well-connected groups of nodes or communities^3,11,12^. In essence, it assesses the segregation of nodes within a network, with higher modularity values indicating a more pronounced community structure. The Louvain method partitions the network into communities in a way that maximizes modularity. The algorithm executes in two primary steps. Initially, each node is designated to its own community. Subsequently, for each node, the algorithm seeks the maximum positive modularity gain by exploring potential movements to all neighboring communities. If no positive gain is achieved, then the node remains in its original community.

The increase of modularity resulting from relocating an isolated node to a community is calculated using the following formula:

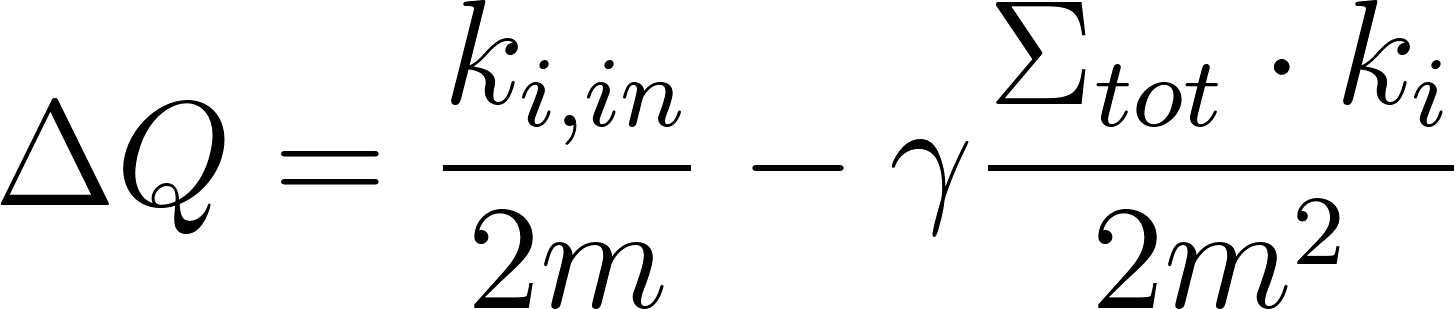
<https://www.codecogs.com/eqnedit.php?latex=%5CDelta%20Q%20%3D%20%5Cfrac%7Bk_%7Bi%2Cin%7D%7D%7B2m%7D%20-%20%5Cgamma%5Cfrac%7B%20%5CSigma_%7Btot%7D%20%5Ccdot%20k_i%7D%7B2m%5E2#0>

Here,
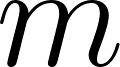
<https://www.codecogs.com/eqnedit.php?latex=m#0> is the graph's size,
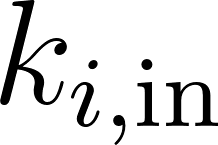
<https://www.codecogs.com/eqnedit.php?latex=k_%7Bi%2C%20%5Ctext%7Bin%7D%7D#0> is the sum of weights of links from the node to nodes in a community,
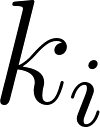
<https://www.codecogs.com/eqnedit.php?latex=k_i#0> is the sum of weights of links incident to the node,
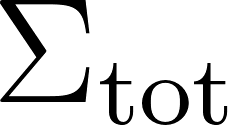
<https://www.codecogs.com/eqnedit.php?latex=%5CSigma_%7B%5Ctext%7Btot%7D%7D#0> is the sum of weights of links incident to nodes in a community, and
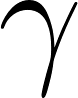
<https://www.codecogs.com/eqnedit.php?latex=%5Cgamma#0> is the resolution parameter.

The first phase iterates until no individual move can enhance the modularity.

In the second phase, a new network is constructed using the communities identified in the first phase as nodes. The weights of links between these new nodes are determined by the sum of weights of links between nodes in the corresponding communities. After completing this phase, the first phase can be reapplied to create larger communities with increased modularity.

Importantly, the number of communities identified by the Louvain method is not fixed or predefined: instead, the algorithm produces molecular communities that arise “naturally” from the input data, without any initial restrictions on the number or sizes of those communities.

##### Identifying the Maximum Weight Spanning Tree

The links that are chosen for inclusion into the spanning tree have the maximum possible combined weight, implying that the respective connected metabolites have the highest similarity scores across the entirety of detected metabolome. In the context of graph theory, a tree is a fundamental structure that consists of nodes connected by edges without forming any cycles. In simpler terms, it represents a connected and acyclic graph. One specific type of tree associated with a graph is a spanning tree. A spanning tree of a graph is a subgraph that includes all the nodes of the original graph while preserving connectivity and remaining acyclic. In computer networking, a spanning tree protocol may be used to ensure a loop-free topology, enabling efficient and reliable communication between network devices^13–16^.

A max-weight spanning tree is a spanning tree where the sum of the weights assigned to its edges is maximized. In other words, it is a tree that spans all the nodes of the graph while emphasizing edges with the highest weights. This concept is particularly relevant in scenarios where the edges of the graph represent weights or costs, and finding a tree that maximizes these weights is essential for certain optimization problems.

Constructing a max-weight spanning tree involves employing algorithms designed for spanning trees, such as Prim's algorithm or Kruskal's algorithm, with a modification to prioritize edges with higher weights. The approach typically involves starting with an arbitrary node, selecting the edge with the maximum weight connected to it, and iteratively adding edges to the tree while avoiding cycles. This process continues until all nodes are included in the tree, resulting in a spanning tree that maximizes the sum of edge weights.

Constructing a maximum weight spanning tree can be achieved using a similar approach as that of finding a minimum weight spanning tree, with a key modification involving the negation of edge weights.

The Kruskal algorithm is a greedy algorithm (algorithm that aims for local optimums at each small stage in order to attempt to reach a global optimum solution) employed for finding the minimum spanning tree (MST) of a connected, weighted graph. The algorithm proceeds through the following steps:

Initialization: Create a forest initially consisting of individual trees, with each tree having a single vertex from the input graph.

Sorting Edges: Sort the edges of the graph in ascending order based on their weights. This step is crucial for the greedy nature of the algorithm, as it considers edges with lower weights first.

Iterative Processing: Loop through the sorted edges. For each edge, perform a cycle test to determine whether adding the edge to the current forest would create a cycle. This is typically done by checking whether the two vertices connected by the edge already belong to the same tree. If adding the edge creates a cycle, skip to the next edge. If not, incorporate the edge into the forest, effectively combining two trees into a single tree.

Termination: Once all edges have been considered, the forest formed by the algorithm constitutes a minimum spanning forest of the graph. If the graph is connected, meaning that all vertices are reachable from any other vertex, the forest will have a single component, forming a minimum spanning tree.

#### Reference Networks of Known Compounds

To corroborate that the appropriate connections are made by MCN, we used reference networks generated for known molecules, both for GC-MS and LC-MS networks. The experimental details are described below. We then explored clustering patterns and evaluated the parameters for network generation and interpretation.

GC-MS. The electron ionization (EI) GC-MS spectra are known to be highly consistent^17^. Therefore, for the reference network, we did not generate reference spectra *de novo*, but instead utilized existing GC-MS spectral reference libraries. The GC-MS reference network shown on Figure S2 was generated by subsetting an open source GC-MS library available at GNPS^18^. The library contains a broad range of compounds that cover various molecular families. A subset of 1000 spectra has been randomly parsed from the library. The molecular community network was generated as described above. The network was then used to explore connectivity patterns of reference compounds (examples of connectivity in different portions of the network are shown in Figures S3-5).

The GNPS job for the unpruned network can be found in Table S1 (Dataset #1).

LC-MS. Since the MS/MS spectra vary for different instruments and modes of acquisition^19^, we have collected reference data on a single instrument, but utilizing several methodologies. This allows excluding the technical variability due to the instrumentation choices, but capturing natural variability of MS/MS spectra for different modes of acquisition. We have collected reference spectra from a library of 800 biologically relevant metabolites purchased from MetaSci (<https://spec.metasci.ca/>). Four different methodologies have been employed to collect LC-MS data from these standards to capture metabolites with different properties: RP, HILIC chromatographies and positive and negative ionization modes. The spectra were then curated and assigned the corresponding annotation provided by the manufacturer. The four libraries were generated and used as an input to generate the LC-MS reference network shown on Figure 1e,f, and S8 as described above. The instrumental settings for each mode of acquisition were as follows.

The GNPS jobs for can be found in Table S1 (Datasets #2-9).

##### Reverse phase (RP) C18 chromatography analysis

For C18 analysis, a Thermo Orbitrap was used in combination with a Waters CSH C18 column (50mm x 2.1 mm x 1.7uM) powered by Vanquish LC system was utilized. Data acquisition was performed using data-dependent acquisition. The gradient elution was performed with mobile phase consisted of 0.1% formic acid in water (Mobile Phase A) and 0.1% formic acid in acetonitrile (Mobile Phase B) with a nominal flow rate of 0.500 ml/min. The autosampler had a draw speed and dispense speed of 100uL/min, and a wash after draw was performed for 3 sec. The samples were maintained at 4°C. The column compartment was set to a temperature of 40°C.

##### HILIC chromatography analysis

The Thermo Orbitrap combined with a BEH-Amide column (50mm x 2.1 mm x 1.7uM) powered by a Vanquish LC system was employed for analysis. Data acquisition was performed using data-dependent acquisition. The gradient elution was performed with mobile phase consisted of 0.01% formic acid, 10mM ammonium formate in water (Mobile Phase A) and 0.1% formic acid, 10mM ammonium formate in acetonitrile (Mobile Phase B) with a nominal flow rate of 0.500 ml/min. The autosampler had a draw speed and dispense speed of 100uL/min, and a wash after draw was performed for 3 sec at a speed of 100 μL/s. Samples were maintained at 4°C. The column compartment was set to a temperature of 40°C.

##### Mass Spectrometry (MS) Parameters C18/HILIC

The application mode was set to Small Molecule, and the method duration was 10 minutes. H-ESI was used as the ion source type, with a static spray voltage of 3500 V for positive ion mode and 2500 V for negative ion mode. The ion transfer tube temperature was set to 350°C, and the vaporizer temperature was set to 400°C. In the MS global settings, infusion mode was selected with an expected LC peak width of 2 seconds. Advanced peak determination was turned on, and default charge state was set to 1. Orbitrap resolution was set at 30000 for MS1 scan and 15000 for MS2 scan, and the scan range mode was set to auto. AGC target was set to standard. Data type was profile, and source fragmentation was disabled. Time mode was set to the retention time window. The mass list table contained compound information including compound name, formula, adduct, m/z, charge state, retention time, window, and polarity for the analytes of interest.

##### MS Data Analysis

The data obtained were converted from the vendor's to mzML format. Feature detection was carried out using Mzmine3^20^, with feature extraction signal threshold set at 5.0E3 and minimum peak width of 2 sec. The mass tolerance was set to 5 ppm, and maximum allowed retention time deviation was set at 10 seconds, based on the statistical evaluation of the quality control samples. For chromatographic deconvolution, the maximum peak width was set at 2 min. Following removal of isotope peaks, the peak lists were aligned using the above-mentioned retention time and mass tolerances. Batches of 50 metabolites were prepared and analyzed with all four different methods. The collected spectra were then searched against existing libraries and the spectra from metabolites that were correctly identified were added to the library. Four separate libraries were generated for each method.

To generate the molecular community network, the compiled libraries were co-networked to generate the unpruned network, followed by the network generation steps described above. We generated six separate community networks with different numbers of matching fragments required to retain an edge between two nodes (Table S1). We then explored the resulting connectivity patterns. Figure 1g, shows the reference network generated with the “Minimum Matched Fragment Ions” setting of 5. The “Organic” layout is used for network visualization.

#### Validation of Novel Bile Acid Conjugates

The biological importance of microbially-derived bile acids has increasingly become prominent since their recent discovery^21–24^. To confirm the structures for the proposed structures shown in Figure 2, we have performed synthesis of candidate compounds at two separate sites, the University of California, San Diego and BieOmix Inc. (Farmington, CT). The results were consistent across the two sites thus enhancing confidence in the structural assignment. The experimental details are described below.

##### Synthesis of postulated conjugates

#### The synthesis of Met-Glu-CA involved sequential activation and coupling of cholic acid with di-tert-butyl L-glutamate and tert-butyl L-methioninate, respectively, followed by selective deprotection of tert-butyl groups, all performed under controlled conditions and subsequently analyzed via LC-MS/MS. Cholic acid was purchased from Sigma Aldrich, di-tert-butyl L-glutamate, and tert-butyl L-methioninate from Chem-Impex Int’l. Inc. HCl-Dioxan 4.0M solution was purchased from Oakwood Chemicals, N,N-Dimethylformamide (DMF) from Merck, 1-Ethyl-3-(3-dimethylaminopropyl)carbodiimide (EDC), N,N-Diisopropylethylamine (DIPEA) from Oakwood Chemicals.

***Glu-CA Synthesis:*** Solid cholic acid (0.12 mmol, 50 mg, 1 eq.) and 2 mL of DMF were added to a 20 mL scintillation vial with a magnetic stir bar. To this solution, solid EDC (0.12 mmol, 23 mg, 1 eq.) and neat DIPEA (0.12 mmol, 21 μl, 1 eq.) were subsequently added, and the solution was stirred at 23°C. After 15 minutes, di-tert-butyl L-glutamate (0.12 mmol, 25 mg, 1 eq.) was added, and the reaction was stirred for 14 hours. The mixture was then concentrated in a vacuum system and used directly to deprotect tert-butyl groups^25,26^. The di-tert-butyl L-glutamate conjugate of cholic acid (0.10 mmol, 65 mg, 1 eq.) and dioxane (1 mL) were added to a 20 mL scintillation vial with a magnetic stir bar. At 0°C, HCl-Dioxane 4.0M solution (1.00 mmol, 0.25 mL, 10 eq.) was added and stirred at room temperature for 2-3 hours. The mixture was then concentrated in a vacuum system and used directly for the next step^26^.

***Met-Glu-CA Synthesis:*** Glu-CA (93 μmol, 50 mg, 1 eq.) and 2 mL of DMF were added to a 20 mL scintillation vial with a magnetic stir bar. To this solution, solid EDC (93 μmol, 18 mg, 1 eq.) and neat DIPEA (93 μmol, 16 μL, 1 eq.) were subsequently added, and the solution was stirred at 23°C. After 15 minutes, tert-butyl L-methioninate (93 μmol, 14 mg, 1 eq.) was added, and the reaction was stirred for 14 hours^25,26^. A tert-butyl L-methioninate conjugate of CA-Glu (80 μmol, 58 mg, 1 eq.) and dioxane (1 ml) were added into a 20 ml scintillation vial with a magnetic stir bar. At 0°C, a 4.0 M HCl-dioxane solution (80 μmol, 20 μl, 10 eq.) was added and stirred at room temperature for 2-3 hours. The mixture was concentrated in a vacuum system and used for LC-MS/MS analysis^27^.

##### Search of novel structures in public data

Note: All the information for the MassIVE datasets mentioned in this section can be found in Supplemental Table S2.

Using the fastMASST^28^ web interface (<https://fasst.gnps2.org/fastsearch/>), we searched for Met-Glu-CA and Glu-Cys-S-S-Cys-CA search against a large-scale metabolomics repository (GNPS/MassIVE). The Universal Spectrum Identifier (USI) of each compound was used as input, for Glu-Cys-S-S-Cys-CA (Spectrum USI: mzspec:GNPS:TASK-4b64ee7363264d5ea92c3aabe3b939a6-spectra/specs_ms.mgf:scan:4162), precursor and fragments mass tolerance were set at 0.05 Da and cosine similarity of 0.7 with at least three matching fragment ions shared between the query spectrum and the available spectra. Glu-Cys-S-S-Cys-CA was found only in a unique dataset that contains 202 strains of human-associated bacteria cultured anaerobically (Spectrum USI: mzspec:MSV000084475:HM_30_BE4_01_45782:scan:1247) (Figure S15).

For Met-Glu-CA (Spectrum USI : mzspec:GNPS2:TASK-90ac45dcf3064aac8fb8f2a9379aa0ac-input_spectra/Alexander_CA_M_G.mzML:scan:2630), precursor and fragments mass tolerance were set at 0.05 Da and cosine similarity of 0.6 with at least three matching fragment ions shared between the query spectrum and the available spectra. This compound was found in six datasets human and microbes related: MSV000081351, MSV000083024, MSV000084218, MSV000084314, MSV000079598, and MSV000080469 (Figure S16, Table S2).

#### Comparison of Community and Conventional Networks

We have explored clustering patterns in conventional and community networks. The molecular community networks for all datasets were generated as described above. We have selected >60 of previously collected datasets available at GNPS/MassIVE^1^ (Table S1, Datasets #10-71). These studies span a variety of sample types, including samples from different uberons of humans, animals, environmental samples, microbial cultures, plants etc., capturing molecular complexity of metabolomics studies. Moreover, we have generated a “global” network of the entirety of metabolome present in the accessible, open data at GNPS/MassIVE^1^, as described below.

##### Global network for the public data available at GNPS

The full network for the publicly available data on GNPS/MassIVE^1^ was generated for LC-MS/MS data as described in^29^. After clustering, a total of 8,453,822 nodes were used to construct the network. Due to such a large number of nodes, an unpruned network would result in an intractably large number of connections, exceeding available computing capabilities; consequently, in order to generate the global network, we applied a top k limit of 10, similarly to conventional networks.

For the MCN, there are 8,040,389 nodes with one or more neighbors. Of the entire graph, 413,433 nodes are singletons and 2,106,277 nodes have zero annotated neighbors (not including the node itself) (Figure 1e-f). When subtracting the nodes with zero annotated neighbors from the nodes with one or more neighbors, there are 5,934,112 nodes with one or more annotated neighbors. For a conventional network with pruning a threshold of 0.7, there are 3,604,775 nodes with one or more neighbors (Figure 1e-f). Of the entire graph, 4,849,047 nodes are singletons and 1,195,998 nodes have zero annotated neighbors (not including the node itself). When subtracting the nodes with zero annotated neighbors from the nodes with one or more neighbors, there are 2,408,777 nodes with one or more annotated neighbors. Therefore, there are approximately additional 3.5 million nodes that become connected to an annotated node in MCN and could be amenable to annotation propagation.

##### Connectivity patterns exploration

The validity of connections in molecular networks requires evaluation to confirm the central hypothesis: that network connections reflect structural relationships between molecules. This assessment is crucial to ensure the network accurately represents molecular structural similarities and can be used for meaningful structural inference and annotation propagation.

For conventional networks, substantial evidence has accumulated since the methodology's introduction over a decade ago, confirming the validity of this central hypothesis^30^. The research community has investigated network connectivity and explored compounds via annotation propagation, followed by structure validation. The pruning threshold of 0.7 emerged empirically from these studies.

Molecular community networks (MCNs) substantially increase connectivity and may introduce connections with cosine scores below the typical similarity threshold of 0.7. Therefore, their validity cannot be automatically assumed based on evidence from conventional networks. The central hypothesis of networking needs empirical investigation for MCNs over a large volume of experimental evidence, similar to conventional networking.

To obtain initial evidence of MCN connection validity, we explored connectivity by elucidating annotations of connected nodes and comparing structures of connected molecules. Our global network encompasses a wide range of research projects with diverse samples, spanning different organisms and biological components, and covering multiple research areas including metabolomics, natural products discovery, environmental analysis, and clinical studies.

As previously described, our global network consisted of 8,453,822 unique nodes. We used this large and diverse dataset to minimize potential biases in molecular distributions that might occur in smaller sample sets, allowing us to explore universal patterns of molecular distribution. To measure structural similarity, we employed the widely established Tanimoto score^31^.

The Tanimoto score was calculated whenever possible, i.e. for all of the edges in the global network where both connected nodes have annotation assigned from library search against 591,778 spectra contained in GNPS community library (https://gnps.ucsd.edu/ProteoSAFe/libraries.jsp). The search identified 1,094,008 annotated nodes out of total 8,453,822 nodes, total of 12.94% of all nodes in the network (overall 2,036,631 nodes had matches with higher than 0.7 cosine similarity, but 942,623 had no SMILES information). We then calculated pairwise Tanimoto scores* for these annotations and compared them for MCN, conventional networks and randomly generated pairs of spectra. The randomly generated pairs were compiled by randomly selecting two features from the same input spectra as for the generated networks, and calculating pairwise cosine scores. For the MCN, Tanimoto similarity could be calculated for 479,608 edges out of 7,963,056 (6.02%). For the conventional network with a pruning threshold of 0.7, Tanimoto similarity could be calculated for 1,618,497 edges out of 17,316,020 edges.

As shown in Figure S17, our results demonstrate that both MCN and conventional networks with a pruning cosine of 0.7 generate non-random connectivity. Both networks produce connections that significantly differ from random pairings and generally capture connections with higher Tanimoto scores at higher cosine values, although this relationship is not direct. In particular, both conventional network and MCN exhibit a large number of edges with high similarity (around cosine 0.7), but low Tanimoto scores (at or below ~0.2). These connections are likely to arise due to at least two factors: possible misannotations and artificially low Tanimoto scores for molecules that differ by multiple substitutions.

Importantly, both networks capture connections with a Tanimoto score of 1, corresponding to nodes with identical annotations. This observation provides crucial validation of the non-random nature of the connectivity. It aligns with the expectation that spectra for the same molecule will be sufficiently similar to be connected in a network, despite variations in MS/MS data across different instruments and experiments.

Furthermore, as a high number of in-source fragments in MS metabolomic data has been predicted^32^, it is corroborated by the observation of a substantial fraction of Tanimoto scores of 1. The GNPS public library used for spectral matching contains spectra for various forms of parent molecules, including different ionization modes and in-source fragmentation products, not just "pure" spectra^1^. Correspondingly, different variants of the same parent structure would contribute to the large number of unity Tanimoto scores, but at a range of cosine similarities, as observed on Figure S17.

These observations support the link between structure and connectivity of both MCN and conventional networks in capturing meaningful molecular relationships, with MCN offering increased connectivity while maintaining structural relevance.

Finally, we show that connectivity in MCN follows a power law distribution (Figure S18). This universal pattern, observed across various complex systems, including biological, social, and technological networks, provides strong evidence that MCN connections are not random but capture fundamental structural patterns within the metabolome. The power law distribution indicates that while most molecules have few connections, some highly connected "hub" molecules play central roles in the network's structure. These hubs likely represent key compounds sharing structural similarities with many other molecules.

*Note: The Tanimoto index was calculated for pairs of SMILES strings using the RDKit library for Python programming language. We used rdFingerprintGenerator with maxPath equal to 7. The code is available at: https://github.com/Alexander0/molecular_communities

#### Supplemental Figures

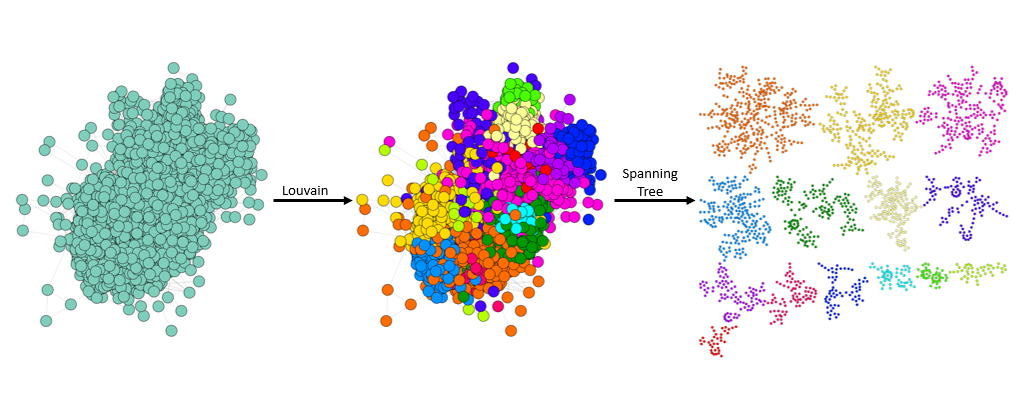

**Figure S1.** Generation of the EI GC-MS spectra network for the GC-MS dataset #18 (Table S1) as described in the “Reference Networks of Known Compounds” section above. The coloring corresponds to the molecular communities detected in the data. From left to right: unpruned network; unpruned network with coloring corresponding to the identified molecular communities; MCN generated by separating sub-networks for communities and pruning them with the Maximum Weight Spanning Tree.

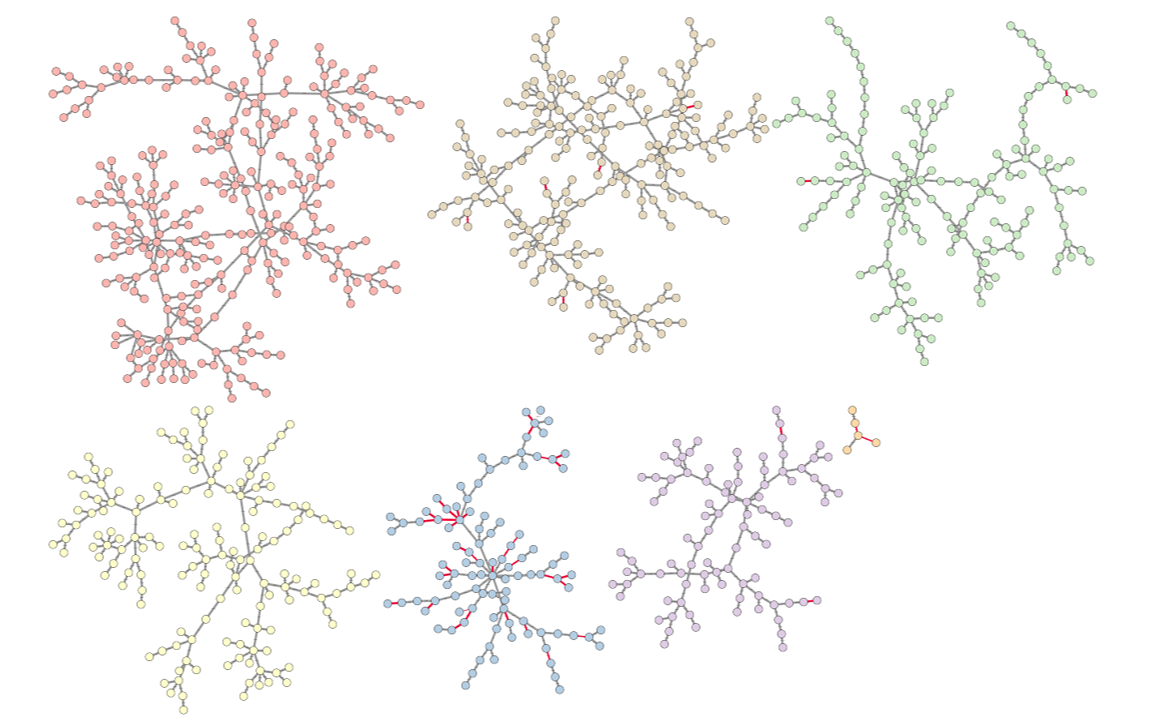

**Figure S2.** A MCN of the reference compounds for EI GC-MS spectra, generated for the subset of 1000 compounds parsed from GNPS library^18^, as described in the “Reference Networks of Known Compounds” section above. The shown network is generated for the dataset #1 (Table S1). The coloring corresponds to the molecular communities detected in the data.

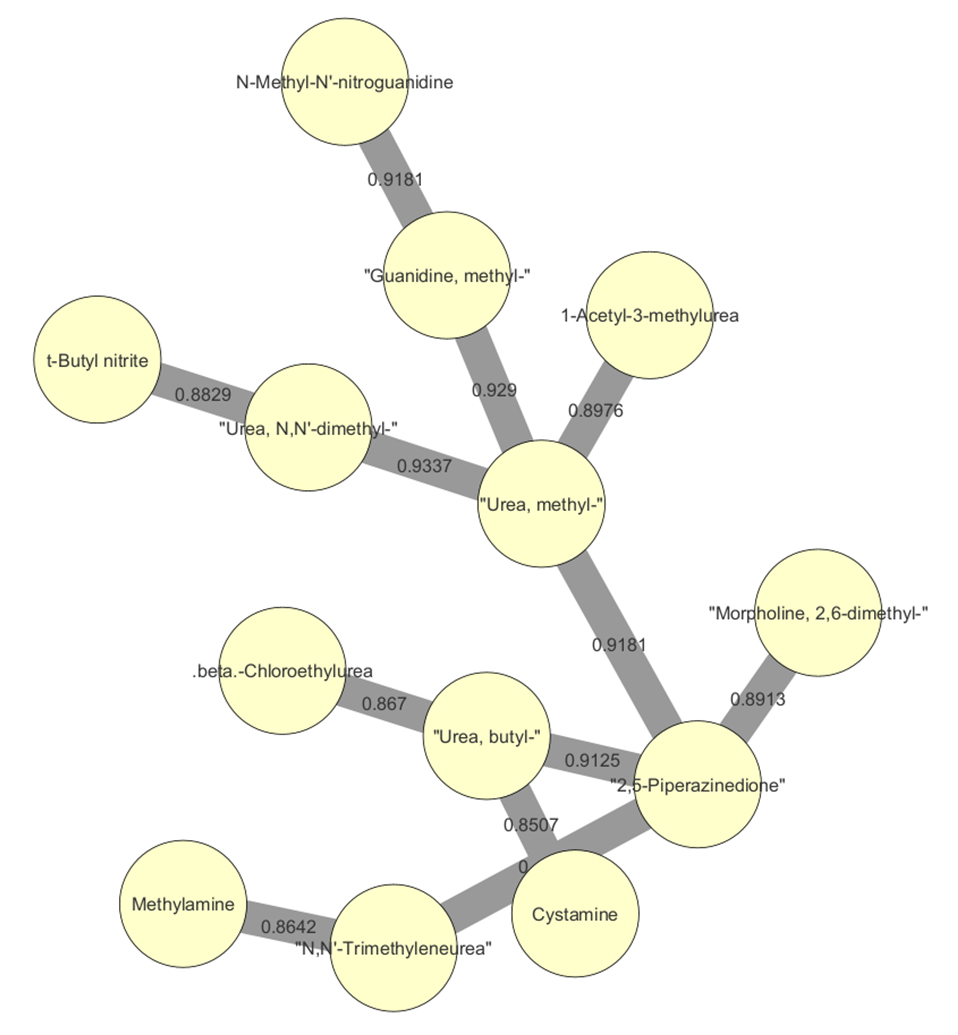

**Figure S3.** An example of MCN connectivity: a close up portion of a molecular community of the MCN shown on Figure S2. The color corresponds to the MCN community. The edge thickness corresponds to the cosine similarity score. A connectivity of amine group-containing compounds is shown.

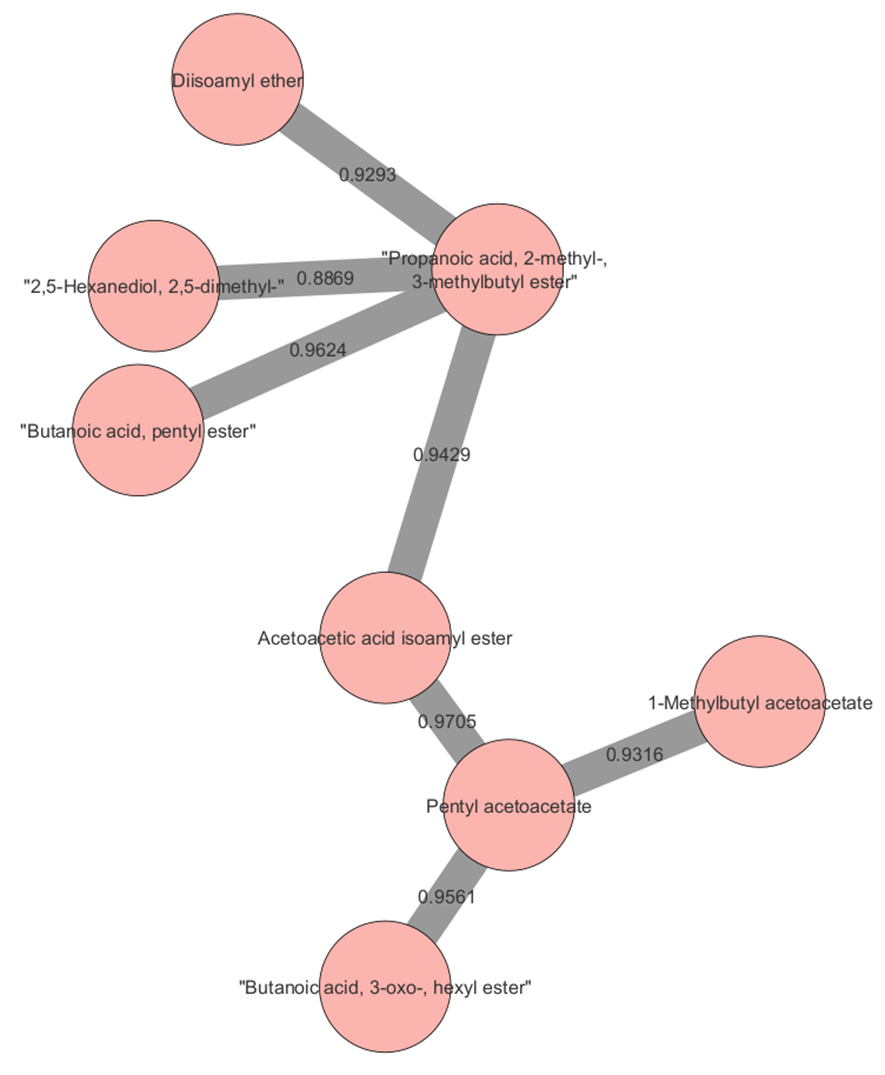

**Figure S4.** An example of MCN connectivity: a close up portion of a molecular community of the MCN shown on Figure S2. The color corresponds to the MCN community. The edge thickness corresponds to the cosine similarity score. A connectivity of esters is shown.

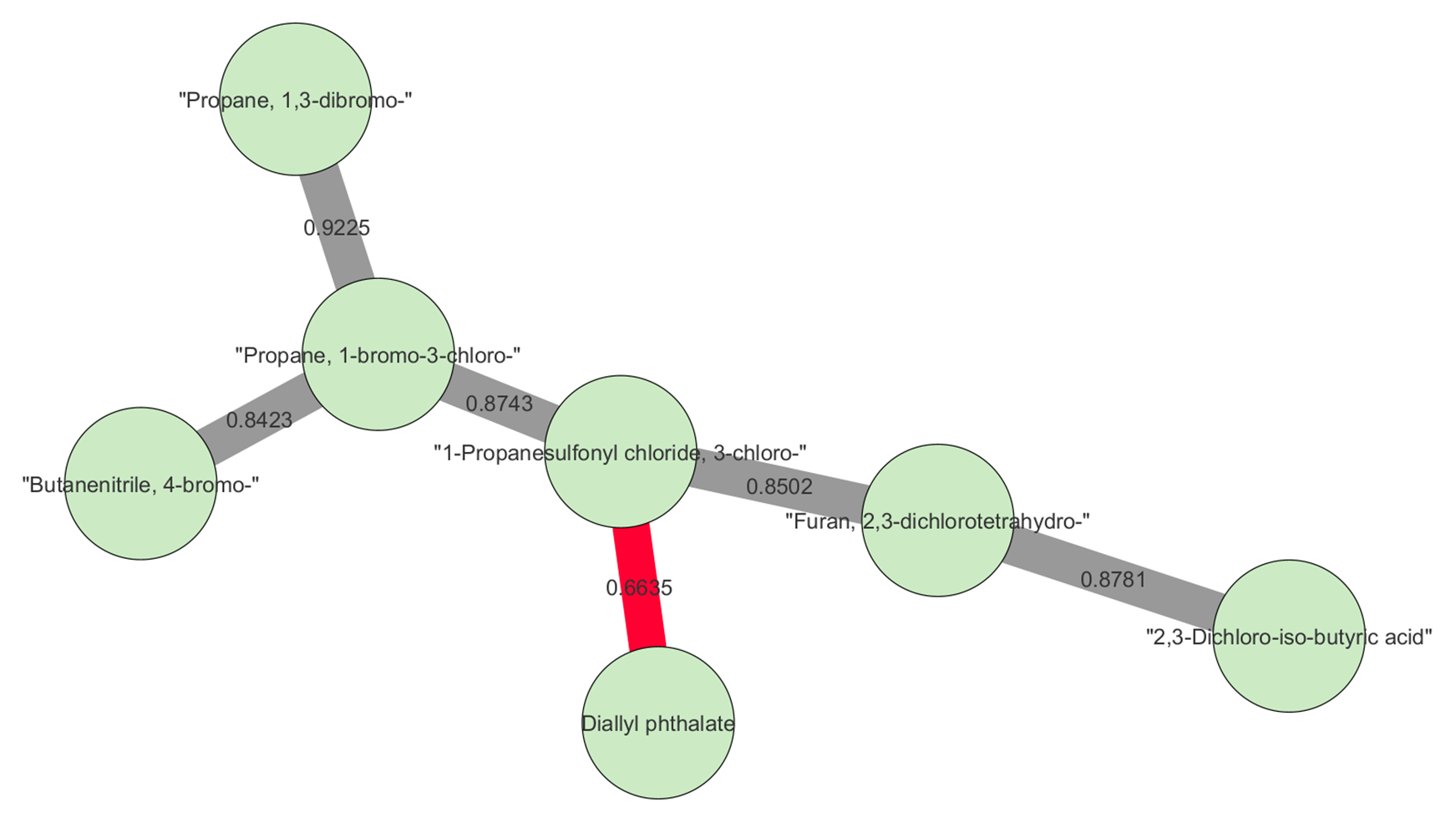

**Figure S5.** An example of MCN connectivity: a close up portion of a molecular community of the MCN shown on Figure S2. The color corresponds to the MCN community; the edge thickness corresponds to the cosine similarity score. The color corresponds to the cosine similarity score: gray, high values (at or above 0.7); red, low values (below 0.7). A connectivity of halogen-containing compounds is shown. This cosine threshold is suggested for high confidence connection between structurally similar nodes in EI GC-MS data.

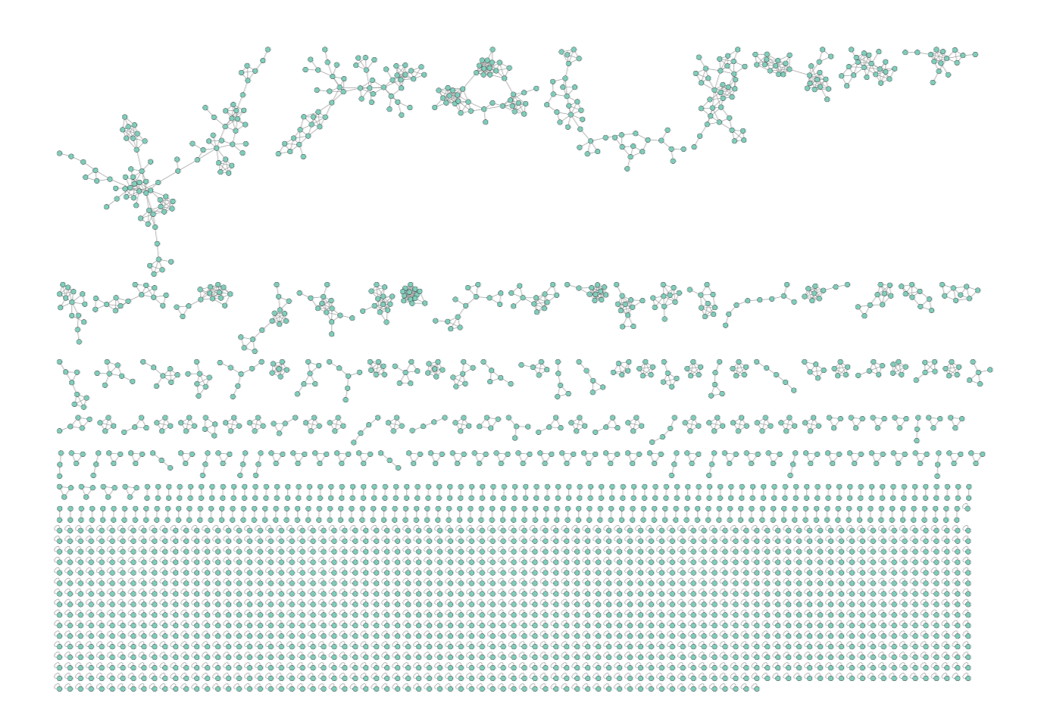

**Figure S6.** A conventional network of the reference compounds for LC-MS/MS spectra, generated for the set of 800 compounds. The pruning cosine similarity threshold is set at 0.7. The shown network is generated for the dataset #2 (Table S1).

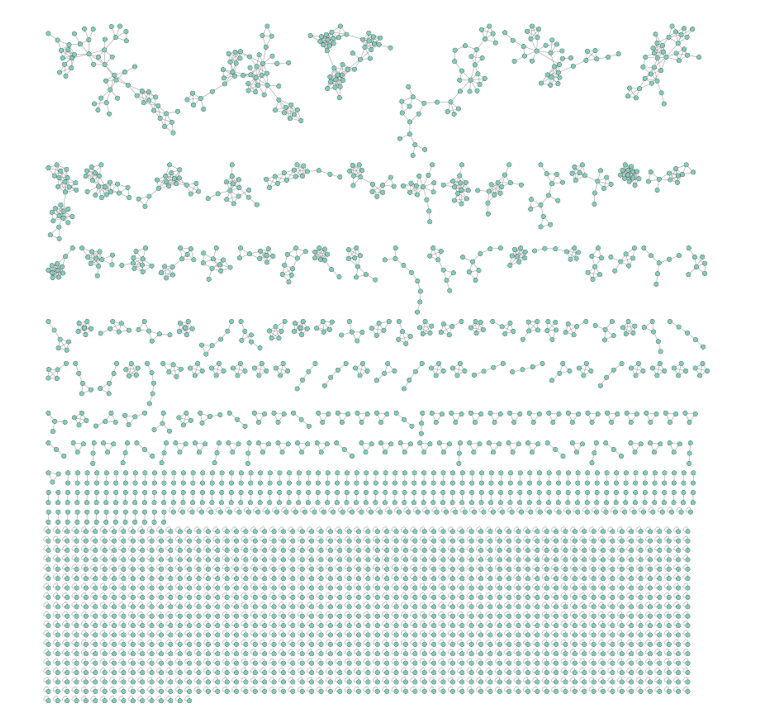

**Figure S7.** A conventional network of the reference compounds for LC-MS/MS spectra, generated for the set of 800 compounds. The pruning cosine similarity threshold is set at 0.5. The shown network is generated for the dataset #3 (Table S1).

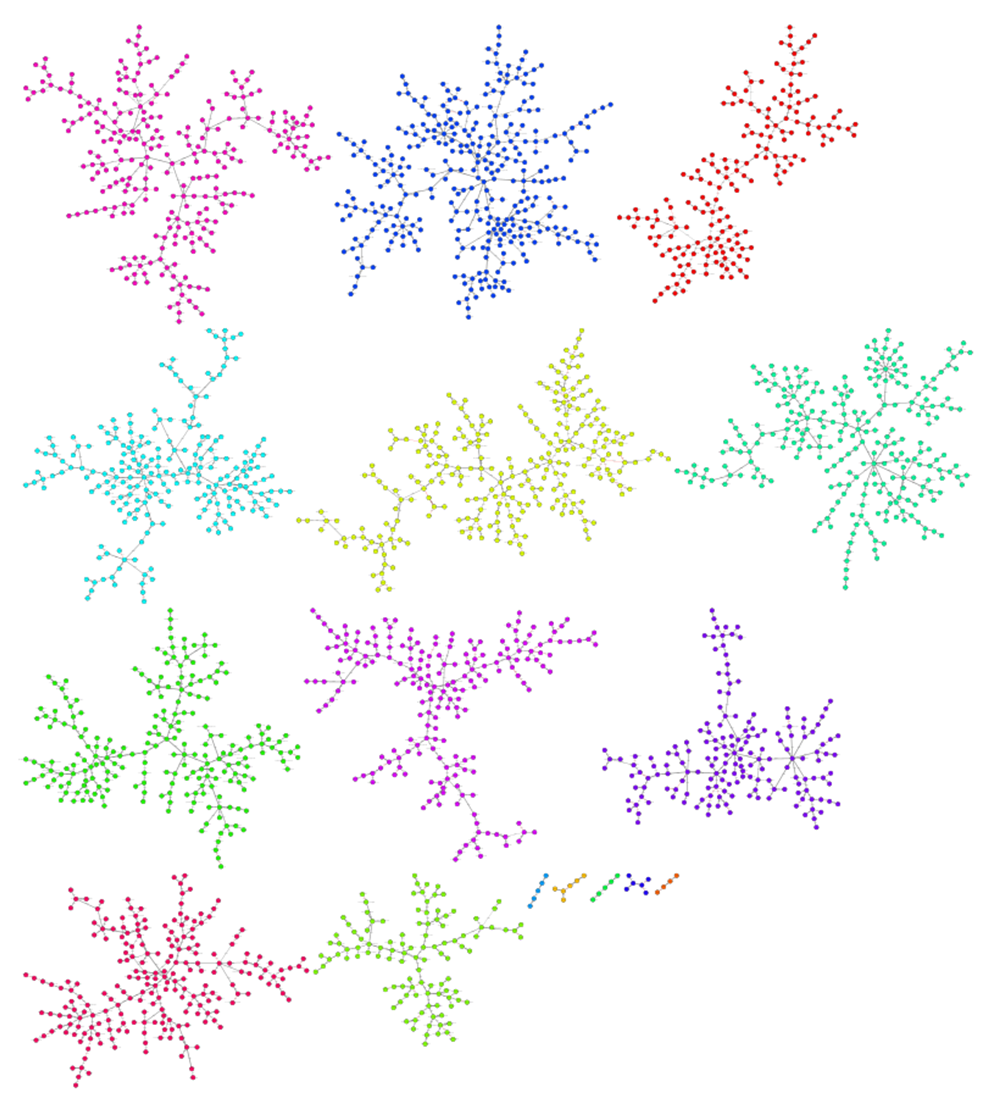

**Figure S8.** A MCN of the reference compounds for LC-MS/MS spectra, generated for the set of 800 compounds, as described in the “Reference Networks of Known Compounds” section above. The shown network is generated for the dataset #7 (Table S1). The coloring corresponds to the molecular communities detected in the data.

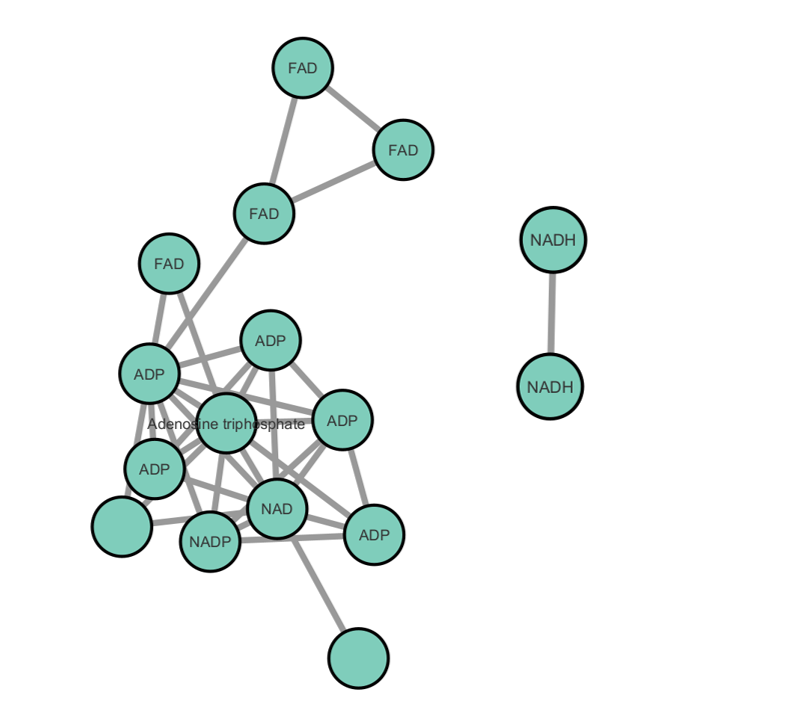

**Figure S9.** A close up of a cluster from the conventional network of the reference compounds shown on Figure S6 (the pruning cosine similarity threshold is set at 0.7). The cluster contains electron carrier metabolites highlighted on Figure 1g.

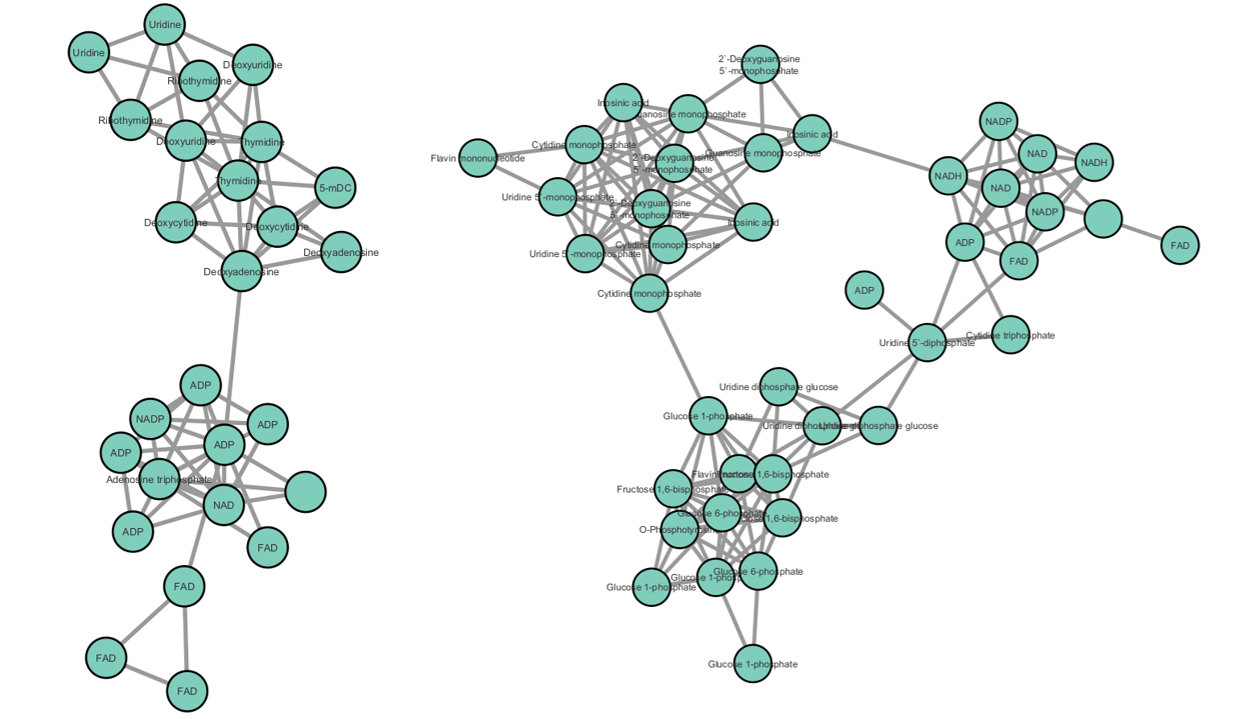

**Figure S10.** A close up of a cluster from the conventional network of the reference compounds shown on Figure S7 (the pruning cosine similarity threshold is set at 0.5). The cluster contains electron carrier metabolites highlighted on Figure 1h.

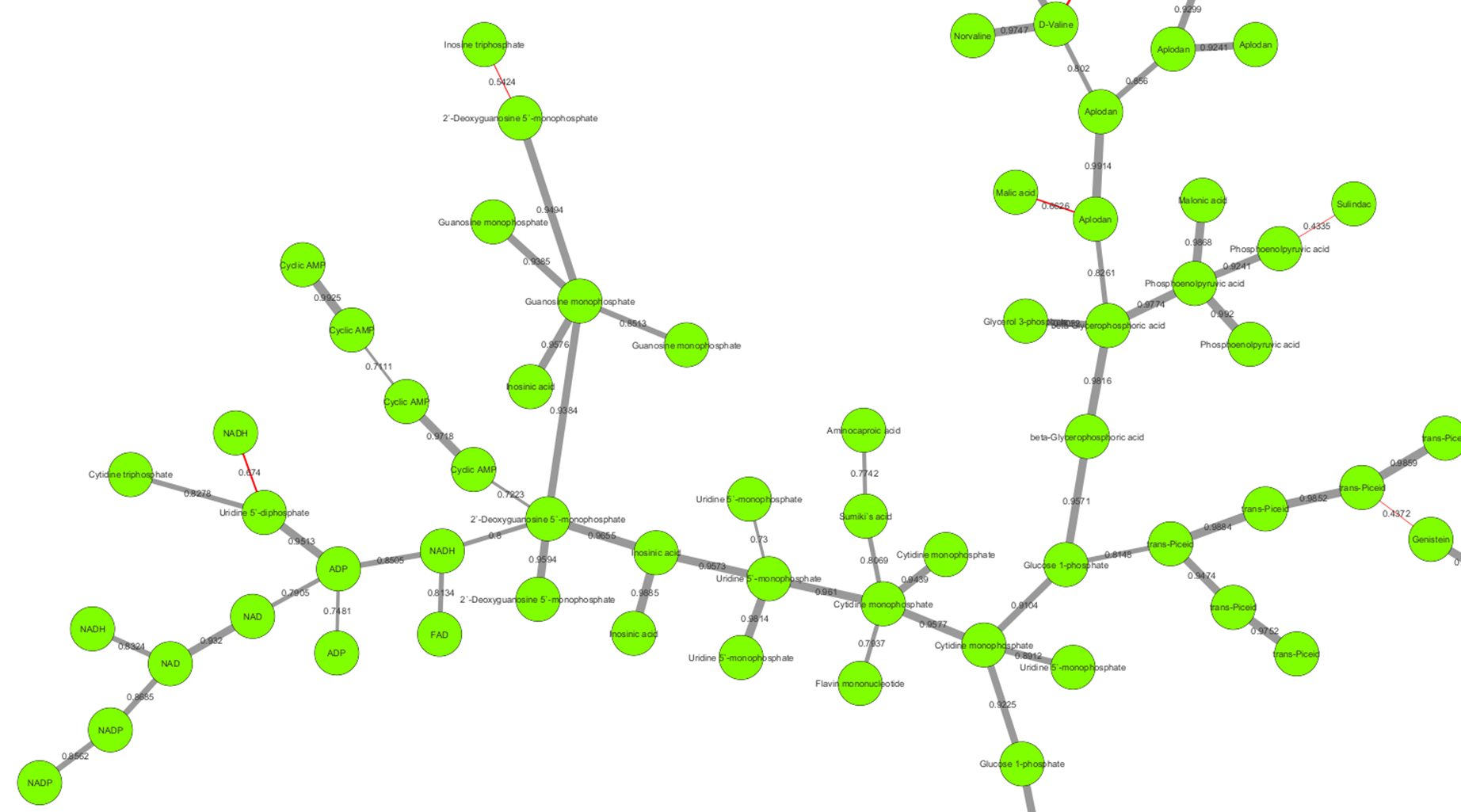

**Figure S11.** A close up of a cluster from MCN of the reference compounds shown on Figure S8. The cluster contains electron carrier metabolites highlighted on Figure 1i.

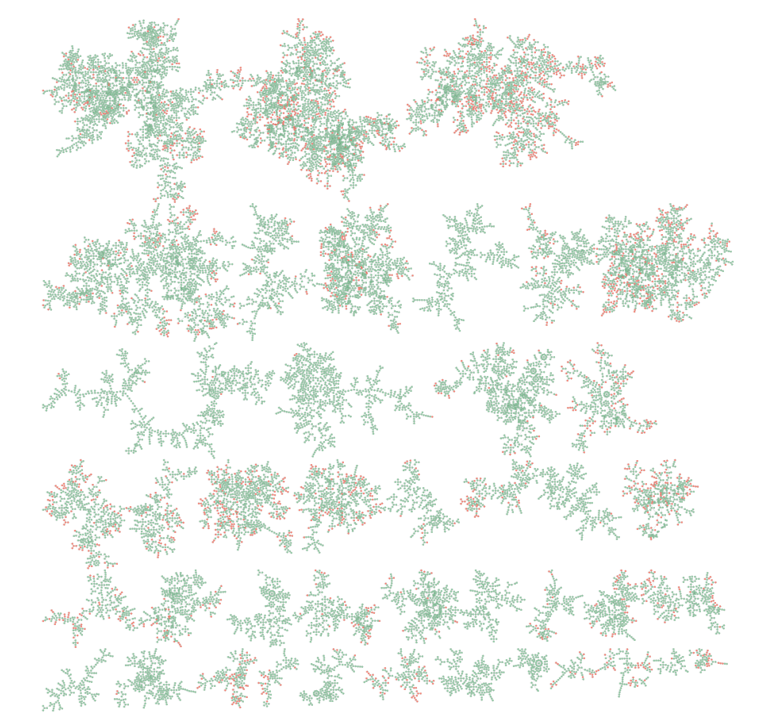

**Figure S12.** MCN generated for the LC-MS/MS data from the American Gut study^33^ (dataset #14 in Table S1). The coloring corresponds to whether the node is connected (green) or disconnected/singleton (red) in the conventional molecular network.

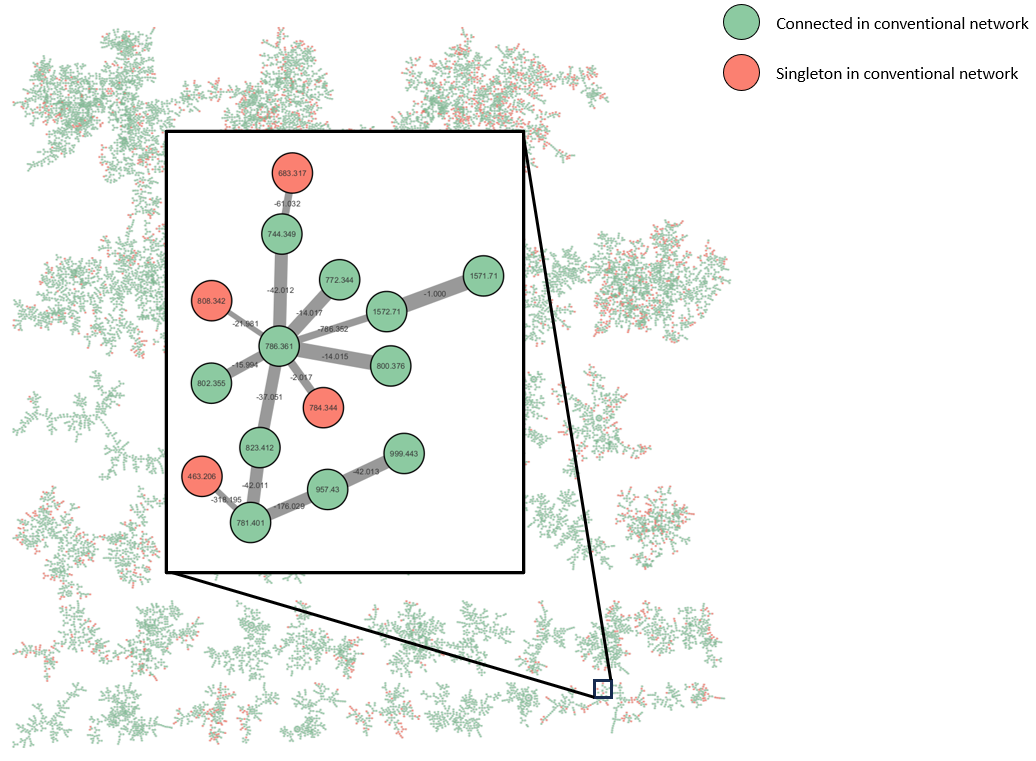

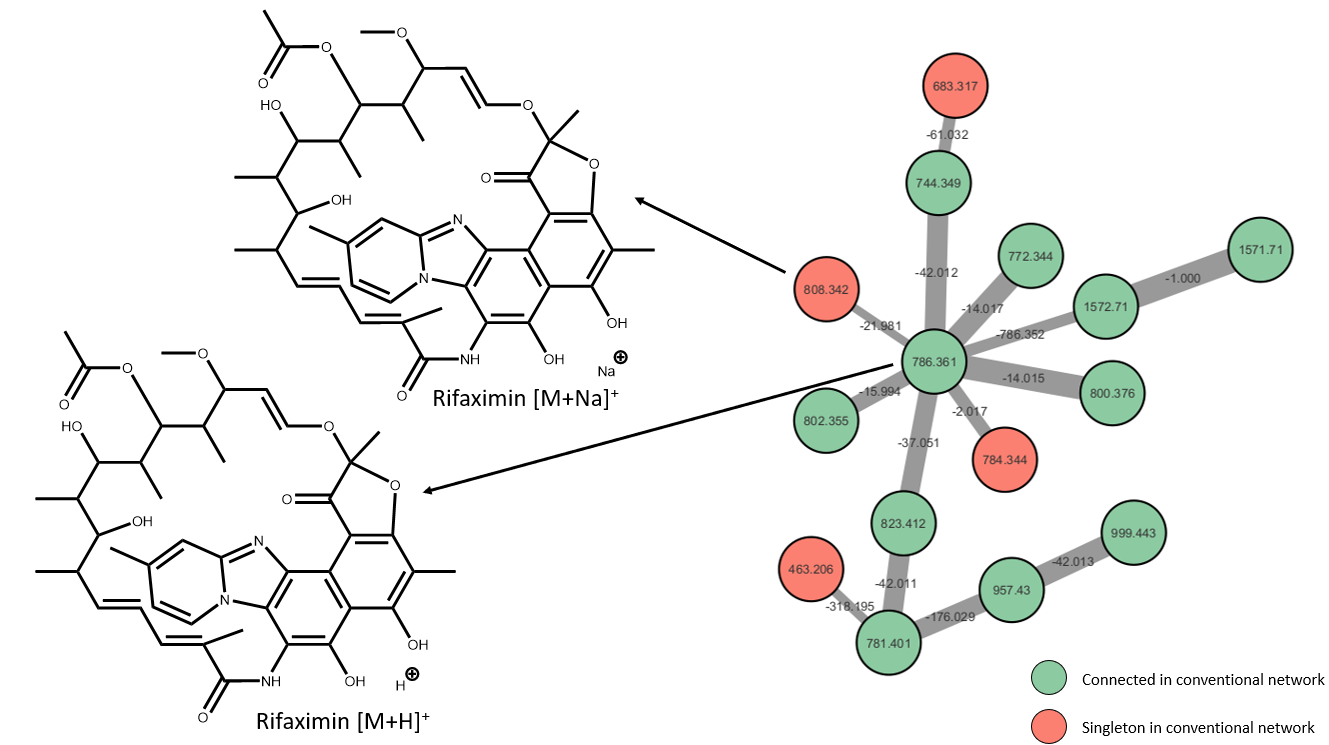

**Figure S13.** The network shown on Figure S11 with a close up of the cluster corresponding to the antibiotic rifaximin, typically used to treat travelers' diarrhea^34^. The coloring corresponds to whether the node is connected (green) or disconnected/singleton (red) in the conventional molecular network; the connection may be in any different clusters across the network. The edge thickness corresponds to the cosine similarity score. Sodiated (and, in general, metal-coordinated) ions exhibit significantly different fragmentation patterns compared to the protonated counterpart. Nodes for such ions would not be connected in conventional networks due to both low cosine score and the correspondingly low probability to be included into top 10 connections, especially for highly connected nodes as in the example of rifaximin. In MCN, the sodiated ion is connected to its parent cluster. Although the similarity of sodiated to protonated rifaximin is relatively low, there is no other molecule more similar to the sodiated rifaximin ion in the data (as is logical and expected), and thus the connection is rescued by MCN.

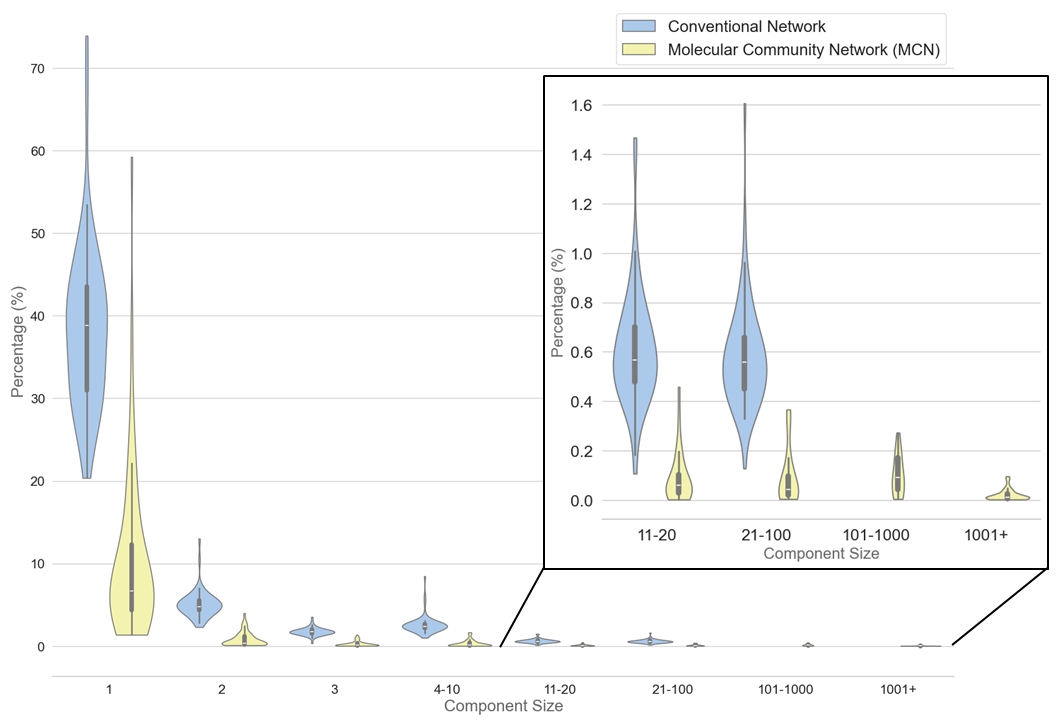

**Figure S14.** Violin plot showing connected components in the conventional (blue) with the pruning cosine threshold of 0.7 and community (yellow) molecular networks for individual datasets listed in the Table S1 (Datasets #11-71). The number of nodes found in clusters with each size k have been counted and the percentage of the nodes within different cluster sizes is shown. Similarly to the global network (Figure 1e-f), in the conventional networks, singletons are predominant. In addition, the larger clusters beyond 100 are not formed due to the artificial cluster size constraints (“Maximum Connected Component Size” setting) needed to maintain limited cluster sizes so the network remains interpretable. Conversely, community networks exhibit on average an order of magnitude lower of singleton and small clusters and naturally form larger clusters. Due to variability of molecular distributions in different datasets, the variability of connectivity is observed.

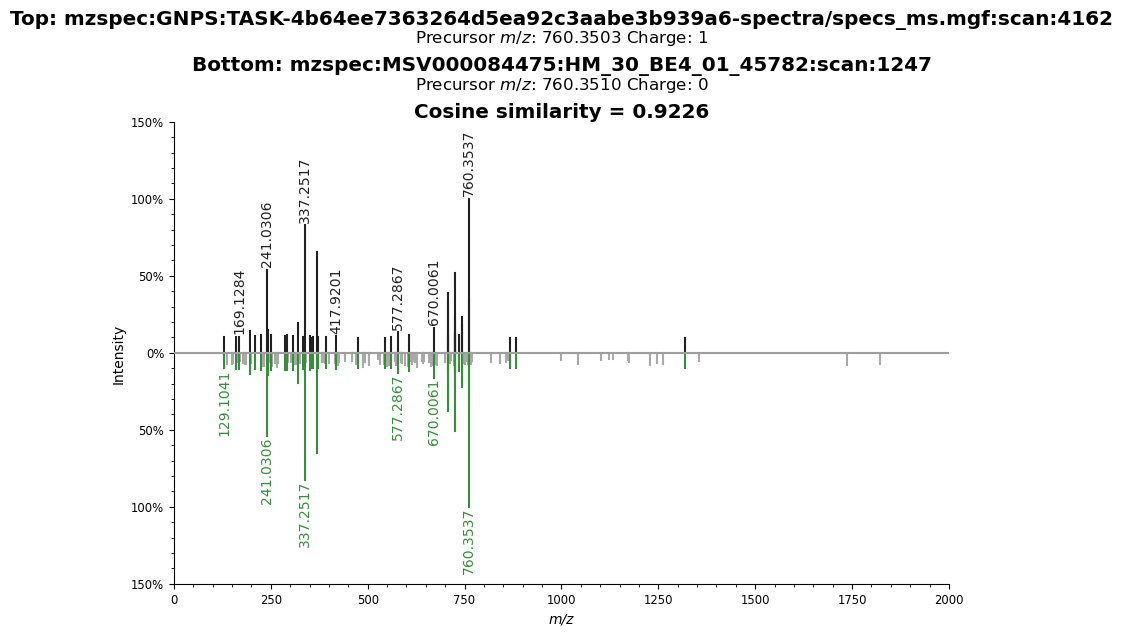

**Figure S15.** Mirror plot between the queried and spectra available in the large-scale metabolomics repository (dataset: MSV000084475).

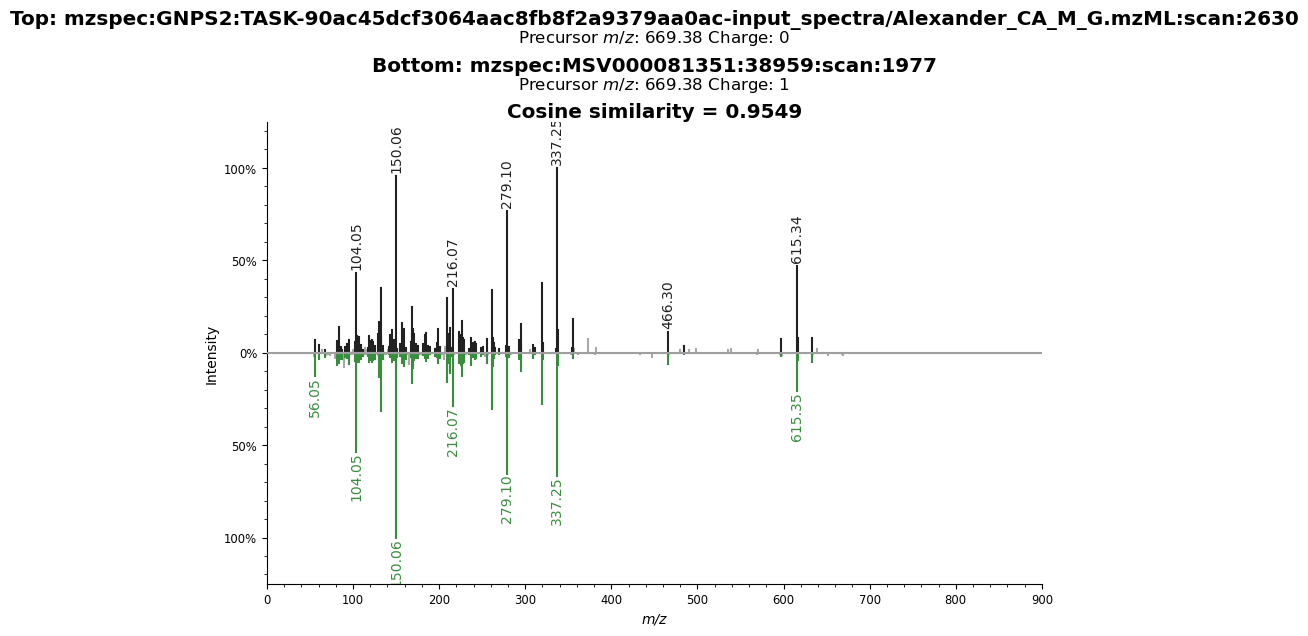

**Figure S16.** Example of mirror plot between the queried and spectra available in the large-scale metabolomics repository (dataset: MSV000081351).

a)
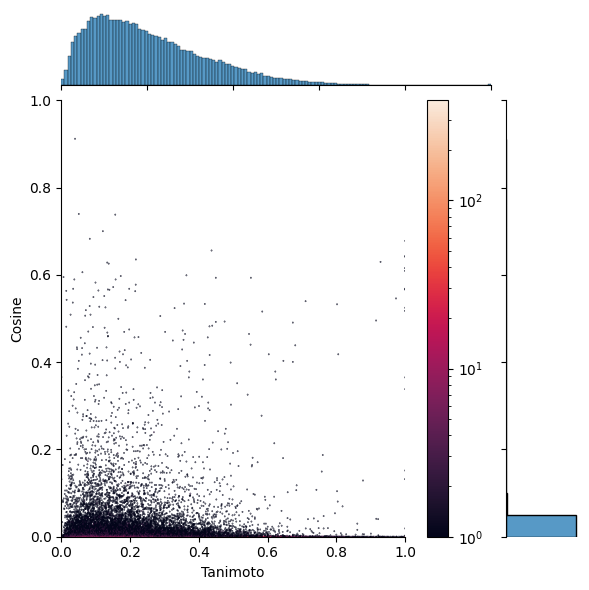

b)
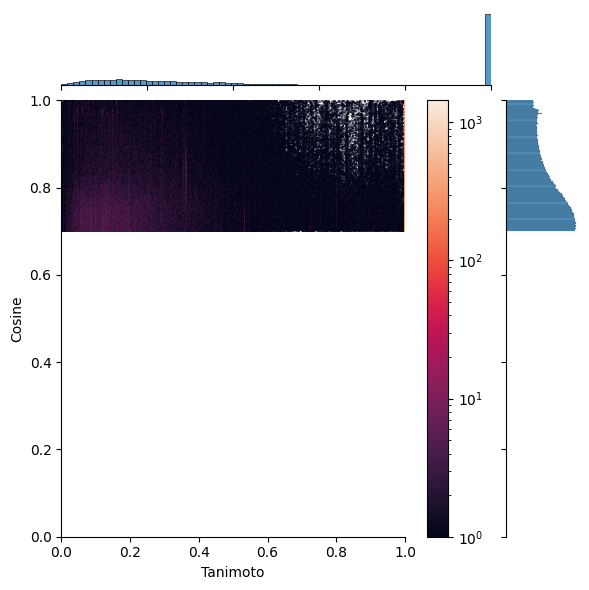

c)
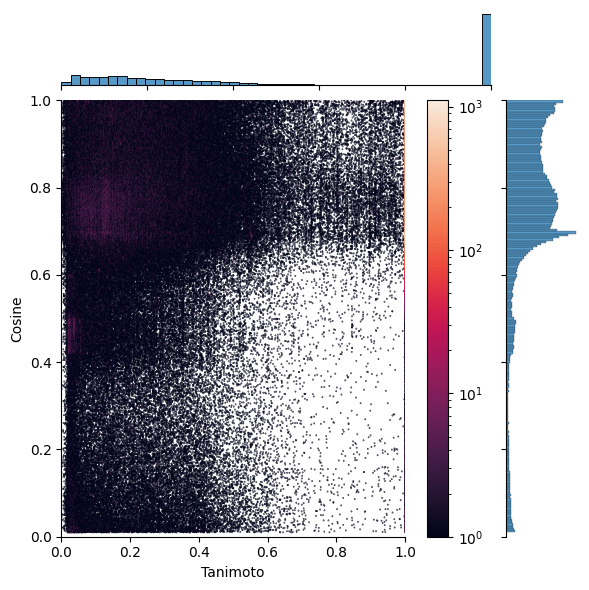

**Figure S17**. Structural similarity analysis in molecular networks assessed for the “global” network generated for the public data available on GNPS, as described in the “Global network for the public data available at GNPS” section. The Tanimoto scores are calculated for linked annotated nodes for: a) randomly selected spectral pairs, b) conventional “global” network with a 0.7 pruning cosine threshold, and c) “global” Molecular Community Networks (MCN). Both MCN and conventional networks show connectivity patterns that clearly differ from random. Beyond the truncation imposed on the conventional network, the connectivity patterns that emerge for MCN and conventional networks appear remarkably similar. The majority of connections in both networks are concentrated at higher cosine values (in MCN, it can be seen that most of connectivity occurs above a cosine similarity of 0.6), indicating a consistent trend in capturing spectral similarities of molecules. Notably, the highest connectivity is shifted towards lower Tanimoto score values (both conventional and community networks display numerous connections with high cosine similarity (~0.7) but low Tanimoto scores (≤0.2)). This shift can be attributed to the inherent limitations of the Tanimoto score as a measure of structural similarity between molecules, particularly for complex structures with multiple modifications, as well as reflecting possible misannotations. The presence of links with Tanimoto scores of 1 in both networks validates their non-random nature, demonstrating successful connection of identically annotated nodes across varying experimental conditions. This also supports the predicted abundance of in-source fragments in MS metabolomic data^32^. Finally, a small number of nodes in MCN are found connected with cosine scores below ~0.3 and exhibit a distribution similar to that of random pairings (shown in panel a), occurring at comparable frequencies. This pattern likely emerges from the presence of noise spectra in the data, (not visible in conventional networks due to pruning). When encountered in MCN, such low-similarity connections could be, therefore, discarded or treated with greater caution.

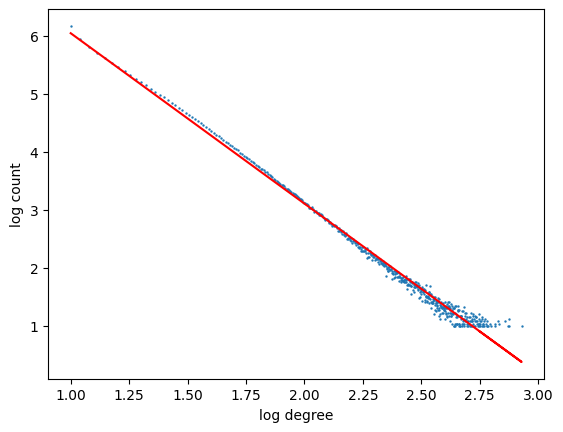

**Figure S18**. Degree distribution of the molecular community network (log-log scale): power-law distribution in the form of N(k) ∝ k^-𝛂^ is observed, where N is the number of nodes with degree k (i.e., connected to k neighbors in the network), and 𝛂 is a positive parameter. In a log-log scale the power-law distribution would be represented by a straight line with a slope of -𝛂. For the plot for molecular communities the linear fit yields 𝛂 = 2.93 (R^2^ = 0.99, for N⩾10 and k⩾10) ). For comparison, the degree distribution of an online social network VK that connects 300M+ users (nodes), was observed to be “bimodal” power-law (piecewise linear in the log-log scale) with 𝛂_1_ = 0.97 (R^2^ = 0.99) for smaller values of k and 𝛂_2_ = 2.28 (R^2^ = 0.98) for larger values of k ^10^. The larger values of the parameter 𝛂 imply faster rates of “decay”, i.e., fewer nodes have large degrees. In the molecular network, a number of links may be missing due to presence of noisy and/or artifactual spectra and irreproducibility of MS/MS patterns across studies, as well as biases of the employed similarity metric, resulting in the increase in value of 𝛂. In contrast, in the online social network all of the links (connections between users) are known. The value of 𝛂 may therefore serve as a metric of the network connectivity “quality” - filling out the molecular space and reduction of lower quality spectra should lead to increased number of links in molecular networks, resulting in corresponding reduction of the value of 𝛂.

### Supplemental Tables

Table S1. Data used in molecular community networking analysis

<https://docs.google.com/spreadsheets/d/1m0UR1yiIVg6Le_11PRDTyiLAHLqLQYr-8ZnDjMYZhnU/edit?usp=sharing>

References cited in Table S1: ^18^^,^^35^^,^^36^^,^^37^^,^^38^^,^^39^

Table S2. Search for novel structures in public data.

<https://docs.google.com/spreadsheets/d/1eF6_JW9a2RF7FgbjhHbah0VpApnJlATqKJvu0eaObWs/edit?usp=sharing>
