## Supplemental Table 1 for "Ordering molecular diversity in untargeted metabolomics via molecular community networking"

| | Dataset description | Methodology | Publication | Massive ID | GNPS link (conventional and/or unpruned network) | MCN properties |
|---|---|---|---|---|---|---|
| 1 | Reference network, GC-MS data | GC-MS | Aksenov, A. A. et al. Auto-deconvolution and molecular networking of gas chromatography-mass spectrometry data. Nat. Biotechnol. 39, 169–173 (2021). | - | https://gnps.ucsd.edu/ProteoSAFe/status.jsp?task=f1648d8e65404aea9ebacac3168b97db | Entire graph: 1000 nodes<br>GCC: 1000 nodes<br>Density: 0.0583<br>Communities: 7<br>Modularity: [illegible] |
| 2 | Reference network (Conventional, cosine threshold 0.7), LC-MS data | LC-MS | - | - | https://gnps.ucsd.edu/ProteoSAFe/status.jsp?task=39c5faf285a549238f3e574eb5176d19 | N/A |
| 3 | Reference network (Conventional, cosine threshold 0.5), LC-MS data | LC-MS | - | - | https://gnps.ucsd.edu/ProteoSAFe/status.jsp?task=6ebe36e1b326434593b8e1812052edb6 | N/A |
| 4 | Reference network, LC-MS data, Minimum Matched Fragment Ions: 1 | LC-MS | - | - | https://gnps.ucsd.edu/ProteoSAFe/status.jsp?task=92625072cea24955904929420944d552 | Entire graph: 2670 nodes<br>GCC: 2456 nodes<br>Density: 0.0156<br>Communities: 16<br>Modularity: 0.5754 |
| 5 | Reference network, LC-MS data, Minimum Matched Fragment Ions: 2 | LC-MS | - | - | https://gnps.ucsd.edu/ProteoSAFe/status.jsp?task=68ae2049bced426896fa9072580c6809 | Entire graph: 2670 nodes<br>GCC: 2432 nodes<br>Density: 0.0154<br>Communities: 14<br>Modularity: 0.5463 |
| 6 | Reference network, LC-MS data, Minimum Matched Fragment Ions: 3 | LC-MS | - | - | https://gnps.ucsd.edu/ProteoSAFe/status.jsp?task=a206b16f9b084429a4e6c96444dfe8d6 | Entire graph: 2670 nodes<br>GCC: 2296 nodes<br>Density: 0.0131<br>Communities: 14<br>Modularity: 0.4669 |
| 7 | Reference network, LC-MS data, Minimum Matched Fragment Ions: 4 | LC-MS | - | - | https://gnps.ucsd.edu/ProteoSAFe/status.jsp?task=c9b26847f8ec4b5ba1a791f1ab40f2b3 | Entire graph: 2670 nodes<br>GCC: 1937 nodes<br>Density: 0.0089<br>Communities: 24<br>Modularity: 0.5521 |
| 8 | Reference network, LC-MS data, Minimum Matched Fragment Ions: 5 | LC-MS | - | - | https://gnps.ucsd.edu/ProteoSAFe/status.jsp?task=efb223d900534ed18765183e03a4a8c | Entire graph: 2670 nodes<br>GCC: 1344 nodes<br>Density: 0.0072<br>Communities: 23<br>Modularity: 0.6858 |
| 9 | Reference network, LC-MS data, Minimum Matched Fragment Ions: 6 | LC-MS | - | - | https://gnps.ucsd.edu/ProteoSAFe/status.jsp?task=33d8913a9a4f4e52b62dc7d9f79c58d8 | Entire graph: 2670 nodes<br>GCC: 794 nodes<br>Density: 0.0089<br>Communities: 21<br>Modularity: 0.7778 |
| 10 | Human Microbiome Project (HMP): human gut microbial cultures biosynthetic potential in bile acid production | LC-MS | Gentry, E. C. et al. Reverse metabolomics for the discovery of chemical structures from humans. Nature 1–3 (2023). | MSV000084475 | https://gnps.ucsd.edu/ProteoSAFe/status.jsp?task=73ede102bc254536aa5cb26ce041556d | Entire graph: 6367 nodes<br>GCC: 5988 nodes<br>Density: 0.0715<br>Communities: 4<br>Modularity: 0.4118 |
| 11 | Volatilome of seaweed | GC-MS | - | - | https://gnps.ucsd.edu/ProteoSAFe/status.jsp?task=57f6118e43d442d3b79ad61a0a7c945e | Entire graph: 1592 nodes<br>GCC: 1586 nodes<br>Density: 0.0275<br>Communities: 14<br>Modularity: 0.7711 |
| 12 | Fecal samples from the study of cirrhosis due to nonalcoholic fatty liver disease | LC-MS | Caussy, C. et al. A gut microbiome signature for cirrhosis due to nonalcoholic fatty liver disease. Nat. Commun. 10, 1–9 (2019). | MSV000082374 | https://gnps.ucsd.edu/ProteoSAFe/status.jsp?task=6b94c41c34ab4aedac4b9e19b40c0c70<br>https://gnps.ucsd.edu/ProteoSAFe/status.jsp?task=6cc7f84b9e4f4cb78e3d9c12f6e6cde9 | Entire graph: 5968 nodes<br>GCC: 4698 nodes<br>Density: 0.0058<br>Communities: 20<br>Modularity: 0.7421 |
| 13 | Volatilome of cheeses | GC-MS | Aksenov, A. A. et al. Auto-deconvolution and molecular networking of gas chromatography-mass spectrometry data. Nat. Biotechnol. 39, 169–173 (2021). | MSV000081161 | https://gnps.ucsd.edu/ProteoSAFe/status.jsp?task=4e52bc2214514aefb19a5de02431897d | Entire graph: 807 nodes<br>GCC: 807 nodes<br>Density: 0.0635<br>Communities: 6<br>Modularity: 0.5897 |
| 14 | American Gut | LC-MS | McDonald, D. et al. American Gut: an Open Platform for Citizen Science Microbiome Research. mSystems 3, (2018) | MSV000080179 | https://gnps.ucsd.edu/ProteoSAFe/status.jsp?task=9d193a46243b47f9bd371788b1bc017d | Entire graph: 25408 nodes<br>GCC: 22053 nodes<br>Density: 0.0018<br>Communities: 44<br>Modularity: 0.8782 |
| 15 | Chemistry of an indoor environment: HomeChem study | LC-MS | Aksenov, A. A. et al. The molecular impact of life in an indoor environment. Science Advances (2022) doi:10.1126/sciadv.abn8016. | MSV000083320 | https://gnps.ucsd.edu/ProteoSAFe/status.jsp?task=679853b1a0df4322926a9617c790f036<br>https://gnps.ucsd.edu/ProteoSAFe/status.jsp?task=066ff333a0eb4854ad111b74f06cd0a3 | Entire graph: 25943 nodes<br>GCC: 23476 nodes<br>Density: 0.0018<br>Communities: 28<br>Modularity: 0.7849 |
| 16 | Murine fecal samples for the study of hepatocarcinoma | LC-MS | Shalapour, S. et al. Inflammation-induced IgA+ cells dismantle anti-liver cancer immunity. Nature 551, 340–345 (2017). | MSV000080918 | https://gnps.ucsd.edu/ProteoSAFe/status.jsp?task=8768ebaf43b04847b77254ffc527abf5<br>https://gnps.ucsd.edu/ProteoSAFe/status.jsp?task=d88f55a8bdc0453fad0a1bc16f0256fd | Entire graph: 6186 nodes<br>GCC: 6027 nodes<br>Density: 0.0063<br>Communities: 37<br>Modularity: 0.7945 |
| 17 | Volatilome of kimchi | GC-MS | - | - | https://gnps.ucsd.edu/ProteoSAFe/status.jsp?task=a19e7f5f59ee45cc981e4fae3f8e878a<br>https://gnps.ucsd.edu/ProteoSAFe/status.jsp?task=d3124aea07bf42ae92eabcd1baeaea52 | Entire graph: 1091 nodes<br>GCC: 1091 nodes<br>Density: 0.0272<br>Communities: 6<br>Modularity: 0.4572 |
| 18 | 3D map of skin volatilome | GC-MS | Aksenov, A. A. et al. Auto-deconvolution and molecular networking of gas chromatography-mass spectrometry data. Nat. Biotechnol. 39, 169–173 (2021). | MSV000084338 | https://gnps.ucsd.edu/ProteoSAFe/status.jsp?task=8aec2d2eb7f64057933fe5ae10a2d731<br>https://gnps.ucsd.edu/ProteoSAFe/status.jsp?task=b0ab41c0d0804599b987f8863d977ff6 | Entire graph: 585 nodes<br>GCC: 513 nodes<br>Density: 0.0098<br>Communities: 21<br>Modularity: 0.8989 |
| 19 | Amino acid-based polymes | LC-MS | - | - | https://gnps.ucsd.edu/ProteoSAFe/status.jsp?task=ef80ef28096845b5af58860d64fef0cb | Entire graph: 1642 nodes<br>GCC: 1431 nodes<br>Density: 0.0395<br>Communities: 10<br>Modularity: 0.7180 |
| 20 | Pseudomonas collection | LC-MS | Nguyen, D. D. et al. Indexing the *Pseudomonas* specialized metabolome enabled the discovery of poaeamide B and the bananamides. Nat. Microbiol. 2, 16197 (2016). | MSV000079450 | https://gnps.ucsd.edu/ProteoSAFe/status.jsp?task=814b9d36b52744f2a87914b47f429673 https://gnps.ucsd.edu/ProteoSAFe/status.jsp?task=22d08e6a6e214d35a31f7ce81bc41901 | Entire graph: 1007 nodes<br>GCC: 858 nodes<br>Density: 0.0250<br>Communities: 14<br>Modularity: 0.6532 |
| 21 | American Gut 3K | LC-MS | - | MSV000080673 | https://gnps.ucsd.edu/ProteoSAFe/status.jsp?task=f12999624bfd4f51a998ab470806e5a2 https://gnps.ucsd.edu/ProteoSAFe/status.jsp?task=adeb80fda38840eabf05a28a27396adf | Entire graph: 22397 nodes<br>GCC: 21254 nodes<br>Density: 0.0021<br>Communities: 49<br>Modularity: 0.9013 |
| 22 | Chemical Standards - Bile Acids & Conjugates, Human | LC-MS | - | MSV000083297 | https://gnps.ucsd.edu/ProteoSAFe/status.jsp?task=3c1d453629b1469581acfae33238d07d https://gnps.ucsd.edu/ProteoSAFe/status.jsp?task=6467b178944a4a29a3572ce9ae2f593e | Entire graph: 606 nodes<br>GCC: 76 nodes<br>Density: 0.1411<br>Communities: 6<br>Modularity: 0.4975 |
| 23 | American Gut Phase 1 | LC-MS | - | MSV000081981 | https://gnps.ucsd.edu/ProteoSAFe/status.jsp?task=348eecfbbdd6424aaea038a40231e224 https://gnps.ucsd.edu/ProteoSAFe/status.jsp?task=aa925ec4ec0840bc9941833f41d0d919 | Entire graph: 20693 nodes<br>GCC: 19698 nodes<br>Density: 0.0023<br>Communities: 43<br>Modularity: 0.8944 |
| 24 | Cyanobacteria Collections | LC-MS | - | MSV000078568 | https://gnps.ucsd.edu/ProteoSAFe/status.jsp?task=ffbfb6c0d5934bdaac6afa9c8a35f69e https://gnps.ucsd.edu/ProteoSAFe/status.jsp?task=5fa41bc64e1749a7a9d3e9ba2ac09cf6 | Entire graph: 21433 nodes<br>GCC: 18336 nodes<br>Density: 0.0014<br>Communities: 54<br>Modularity: 0.8811 |

| | | | | | | |
|---|---|---|---|---|---|---|
| 25 | Undiagnosed Disease Network - Plasma Lipidomics and Metabolomics | GC-MS (metabolomics) | Kyle, JE. et al. A resource of lipidomics and metabolomics data from individuals with undiagosed diseases. Sci Data 8(1): 114. (2021). | MSV000084716 | https://gnps.ucsd.edu/ProteoSAFe/status.jsp?task=c22152591cf94f3aa0e5171efbede16d https://gnps.ucsd.edu/ProteoSAFe/status.jsp?task=a32ff415d15d4e03ba5b0b489b05acaa https://gnps.ucsd.edu/ProteoSAFe/status.jsp?task=ff0536be855246aeb5227ed287735129 | Entire graph: 517 nodes<br>GCC: 517 nodes<br>Density: 0.0750<br>Communities: 7<br>Modularity: 0.4358 |
| 26 | Undiagnosed Disease Network - Plasma Lipidomics and Metabolomics | Negative Mode LC-MS (lipidomics) | Kyle, JE. et al. A resource of lipidomics and metabolomics data from individuals with undiagosed diseases. Sci Data 8(1): 114. (2021). | MSV000084716 | https://gnps.ucsd.edu/ProteoSAFe/status.jsp?task=3c3da031c17f4ba09d1af597de0fa110 https://gnps.ucsd.edu/ProteoSAFe/status.jsp?task=432b8d7d561d4e2d841495c45e017d34 | Entire graph: 12513 nodes<br>GCC: 6667 nodes<br>Density: 0.0025<br>Communities: 21<br>Modularity: 0.7652 |
| 27 | Undiagnosed Disease Network - Plasma Lipidomics and Metabolomics | Positive Mode LC-MS (lipidomics) | Kyle, JE. et al. A resource of lipidomics and metabolomics data from individuals with undiagosed diseases. Sci Data 8(1): 114. (2021). | MSV000084716 | https://gnps.ucsd.edu/ProteoSAFe/status.jsp?task=18a45d74397f4ff1ac890521832dc0ee https://gnps.ucsd.edu/ProteoSAFe/status.jsp?task=7ace8dbca4304e6a9b16493fad7f2dc8 | Entire graph: 21117 nodes<br>GCC: 15521 nodes<br>Density: 0.0011<br>Communities: 40<br>Modularity: 0.8906 |
| 28 | Multi-Omics Analysis of COVID-19 Severity | GC-MS (metabolomics) | Overmyer, KA. et al. Large-scale Multi-omic Analysis of COVID-19 Severity. Cell Syst. 12(1): 23 - 40. (2021) | MSV000085703 | https://gnps.ucsd.edu/ProteoSAFe/status.jsp?task=72de42710cde419aa084364c3fa8dd49 https://gnps.ucsd.edu/ProteoSAFe/status.jsp?task=9a71d35a4672480180274a8f419f65fb https://gnps.ucsd.edu/ProteoSAFe/status.jsp?task=3c8cfe35da224bbdb58657633b22a19d | Entire graph: 514 nodes<br>GCC: 514 nodes<br>Density: 0.0812<br>Communities: 5<br>Modularity: 0.4084 |
| 29 | Multi-Omics Analysis of COVID-19 Severity | LC-MS (lipidomics) | Overmyer, KA. et al. Large-scale Multi-omic Analysis of COVID-19 Severity. Cell Syst. 12(1): 23 - 40. (2021) | MSV000085703 | https://gnps.ucsd.edu/ProteoSAFe/status.jsp?task=dc6271823a13456b887158d371bd8831 https://gnps.ucsd.edu/ProteoSAFe/status.jsp?task=d7a1d6a7b7a141a09eb9551add953e97 | Entire graph: 7477 nodes<br>GCC: 7042 nodes<br>Density: 0.0056<br>Communities: 30<br>Modularity: 0.8726 |
| 30 | Multi-Omics Analysis of COVID-19 Severity | LC-MS (proteomics) | Overmyer, KA. et al. Large-scale Multi-omic Analysis of COVID-19 Severity. Cell Syst. 12(1): 23 - 40. (2021) | MSV000085703 | https://gnps.ucsd.edu/ProteoSAFe/status.jsp?task=c1947bd6b8e543a19bcc2bf0af717d6a https://gnps.ucsd.edu/ProteoSAFe/status.jsp?task=a2d39c1cde5f44d4899fc6e3668b2d05 | Entire graph: 2091 nodes<br>GCC: 971 nodes<br>Density: 0.0544<br>Communities: 9<br>Modularity: 0.6699 |
| 31 | Extracts of HMP Project Isolates | LC-MS | - | MSV000082045 | https://gnps.ucsd.edu/ProteoSAFe/status.jsp?task=e2850a91beb2461aa00a8743b8a41a4a https://gnps.ucsd.edu/ProteoSAFe/status.jsp?task=cefb818e205744868aa5d6c33bacd5c6 | Entire graph: 8461 nodes<br>GCC: 7869 nodes<br>Density: 0.0050<br>Communities: 24<br>Modularity: 0.7618 |
| 32 | Leaf Tissues of Orange Trees | LC-MS | - | MSV000082963 | https://gnps.ucsd.edu/ProteoSAFe/status.jsp?task=cf8d28c84d5e425697b9feb4cac729c5 https://gnps.ucsd.edu/ProteoSAFe/status.jsp?task=cb5556a1c175406783bdcda8b2d3ea88 | Entire graph: 7324 nodes<br>GCC: 6879 nodes<br>Density: 0.0043<br>Communities: 27<br>Modularity: 0.7510 |
| 33 | Human Blood Serum Spiked with Standards of FAMEs and Hydrocarbons | GC-MS | - | MSV000085128 | https://gnps.ucsd.edu/ProteoSAFe/status.jsp?task=52a8bdbac4c648ad9a501b48e3b22af7 https://gnps.ucsd.edu/ProteoSAFe/status.jsp?task=19ac8e9e73f841b18a457ac4f65fd59e https://gnps.ucsd.edu/ProteoSAFe/status.jsp?task=f542a5565a3b48e09b3b8e846fa6b820 | Entire graph: 367 nodes<br>GCC: 367 nodes<br>Density: 0.1560<br>Communities: 5<br>Modularity: 0.2778 |
| 34 | Skin Microbiome | LC-MS | - | MSV000081582 | https://gnps.ucsd.edu/ProteoSAFe/status.jsp?task=b28ab360b34c4066ba961c759afc6a52 https://gnps.ucsd.edu/ProteoSAFe/status.jsp?task=15ecdb191abc4f67996c9fe2cf253e9c | Entire graph: 18135 nodes<br>GCC: 15843 nodes<br>Density: 0.0028<br>Communities: 44<br>Modularity: 0.9237 |
| 35 | Global Foodomics | LC-MS | - | MSV000084900 | https://gnps.ucsd.edu/ProteoSAFe/status.jsp?task=e93d7d2240924d78b66da490bce42172 https://gnps.ucsd.edu/ProteoSAFe/status.jsp?task=5b2eb5fe716c4d63ada84f46ad526b79 | Entire graph: 43545 nodes<br>GCC: 41972 nodes<br>Density: 0.0009<br>Communities: 48<br>Modularity: 0.8244 |
| 36 | Animal Species Bile Extracts | LC-MS | - | MSV000085120 | https://gnps.ucsd.edu/ProteoSAFe/status.jsp?task=f80a9dcc73fd476da4db995b1e0ff253 https://gnps.ucsd.edu/ProteoSAFe/status.jsp?task=243724c318504f1a9174889094b131dc | Entire graph: 32695 nodes<br>GCC: 31750 nodes<br>Density: 0.0013<br>Communities: 38<br>Modularity: 0.8698 |
| 37 | Human Milk Samples | LC-MS | - | MSV000091520 | https://gnps.ucsd.edu/ProteoSAFe/status.jsp?task=ea82e15cda88453e9777bfc55c99ba91 https://gnps.ucsd.edu/ProteoSAFe/status.jsp?task=cec94dc7b060472d8830cb8f622ccd6e | Entire graph: 26206 nodes<br>GCC: 25335 nodes<br>Density: 0.0019<br>Communities: 38<br>Modularity: 0.8925 |
| 38 | Sputum Samples from Cystic Fibrosis Patients | LC-MS | - | MSV000082667 | https://gnps.ucsd.edu/ProteoSAFe/status.jsp?task=1be0e32a09fe4eab80dc3133daff6d37 https://gnps.ucsd.edu/ProteoSAFe/status.jsp?task=9cc04dea68d0463c8c22db6bff786383 | Entire graph: 13830 nodes<br>GCC: 12762 nodes<br>Density: 0.0027<br>Communities: 51<br>Modularity: 0.8572 |
| 39 | Swabs of Hands and Objects | LC-MS | - | MSV000080030 | https://gnps.ucsd.edu/ProteoSAFe/status.jsp?task=ac60ddffccda4611887d3ff391f97bf5 https://gnps.ucsd.edu/ProteoSAFe/status.jsp?task=d4a33b56c6ef4a5f83d11dc90e4d2635 | Entire graph: 17655 nodes<br>GCC: 15527 nodes<br>Density: 0.0025<br>Communities: 47<br>Modularity: 0.8961 |
| 40 | Metabolome from Organs of Germ Free & Pathogen Free Mice | LC-MS | - | MSV000079949 | https://gnps.ucsd.edu/ProteoSAFe/status.jsp?task=6c5f40224da346ccbaf0437e846e3c48 https://gnps.ucsd.edu/ProteoSAFe/status.jsp?task=eada7d7bef8b46faa308412ea8ffb506 | Entire graph: 9730 nodes<br>GCC: 7653 nodes<br>Density: 0.0039<br>Communities: 46<br>Modularity: 0.8875 |
| 41 | Microbial Cultures | LC-MS | - | MSV000090268 | https://gnps.ucsd.edu/ProteoSAFe/status.jsp?task=de263e73196d4254bbd571aeb765cee0 https://gnps.ucsd.edu/ProteoSAFe/status.jsp?task=9db43585ebc8432a911d5894622931e7 | Entire graph: 23805 nodes<br>GCC: 15877 nodes<br>Density: 0.0019<br>Communities: 62<br>Modularity: 0.8769 |
| 42 | Human Teeth Powders | LC-MS | - | MSV000082869 | https://gnps.ucsd.edu/ProteoSAFe/status.jsp?task=93345ed3dcfa4db69c38aa0e1c9528ac https://gnps.ucsd.edu/ProteoSAFe/status.jsp?task=cbd2501cabaf43f5ae5b9899e8e55741 | Entire graph: 9540 nodes<br>GCC: 8814 nodes<br>Density: 0.0038<br>Communities: 21<br>Modularity: 0.7555 |
| 43 | Metabolomic Samples from Brain Cohort | LC-MS | - | MSV000086415 | https://gnps.ucsd.edu/ProteoSAFe/status.jsp?task=efefaf28d4b146b4b2176effe7936d0c https://gnps.ucsd.edu/ProteoSAFe/status.jsp?task=15b8944c23964f59b4c2dfbc1e99dea4 | Entire graph: 10825 nodes<br>GCC: 10186 nodes<br>Density: 0.0037<br>Communities: 29<br>Modularity: 0.8177 |
| 44 | Metabolomic Samples from Plasma Cohort | LC-MS | - | MSV000086270 | https://gnps.ucsd.edu/ProteoSAFe/status.jsp?task=da133c713c4f4a8cb5fb2c266dc83d6e https://gnps.ucsd.edu/ProteoSAFe/status.jsp?task=d0f73619fc2845618f373d3679b9e683 | Entire graph: 7334 nodes<br>GCC: 6839 nodes<br>Density: 0.0060<br>Communities: 26<br>Modularity: 0.8455 |
| 45 | Human Skin Microbiome Isolates | LC-MS | Timm, CM., et al. Isolation and characterization of diverse microbal representatives from the human skin microbiome. Microbiome 8(58). (2020). | MSV000086550 | https://gnps.ucsd.edu/ProteoSAFe/status.jsp?task=f29754bb10db4f9aa4e9cdce175c3ca0 https://gnps.ucsd.edu/ProteoSAFe/status.jsp?task=3f1dbc7e141142a58d3e201569770417 | Entire graph: 8374 nodes<br>GCC: 8051 nodes<br>Density: 0.0054<br>Communities: 30<br>Modularity: 0.8423 |
| 46 | Drug Metabolism Study | LC-MS | - | MSV000082493 | https://gnps.ucsd.edu/ProteoSAFe/status.jsp?task=9bbc7ad76f71439ba85f6a73bb326e52 https://gnps.ucsd.edu/ProteoSAFe/status.jsp?task=136d77d0a95b4c9da972a00b6948e72c | Entire graph: 8620 nodes<br>GCC: 8155 nodes<br>Density: 0.0048<br>Communities: 31<br>Modularity: 0.8335 |
| 47 | Amazon Skin and Environment | LC-MS | - | MSV000079389 | https://gnps.ucsd.edu/ProteoSAFe/status.jsp?task=c44891c85fe540d79d2aa3173bd5570d https://gnps.ucsd.edu/ProteoSAFe/status.jsp?task=a3a691e8a271426799ada6adcd669814 | Entire graph: 14420 nodes<br>GCC: 12343 nodes<br>Density: 0.0035<br>Communities: 33<br>Modularity: 0.8298 |
| 48 | Waimea Supertransect Metabolomics | LC-MS | - | MSV000085129 | https://gnps.ucsd.edu/ProteoSAFe/status.jsp?task=3963175ca98c4409b68ce0c1328f538e https://gnps.ucsd.edu/ProteoSAFe/status.jsp?task=219727a724d74b91af25b82813aab118 | Entire graph: 9655 nodes<br>GCC: 8768 nodes<br>Density: 0.0042<br>Communities: 26<br>Modularity: 0.7252 |

| | | | | | | |
|---|---|---|---|---|---|---|
| 49 | Actinobacteria | LC-MS | - | MSV000078688 | https://gnps.ucsd.edu/ProteoSAFe/status.jsp?task=4bab86adbbb24e3dbc99a87cc2743f1f https://gnps.ucsd.edu/ProteoSAFe/status.jsp?task=e1f8e9ac265140b597f0af6c4c2a39e6 | Entire graph: 1092 nodes<br>GCC: 568 nodes<br>Density: 0.0233<br>Communities: 19<br>Modularity: 0.8381 |
| 50 | Venezuela Urbanization Gradient | LC-MS | - | MSV000082433 | https://gnps.ucsd.edu/ProteoSAFe/status.jsp?task=c22b8b97333149618c7c5f56e521bb9b https://gnps.ucsd.edu/ProteoSAFe/status.jsp?task=aec7743f8db94eacae865cda4441c39a | Entire graph: 14674 nodes<br>GCC: 14182 nodes<br>Density: 0.0029<br>Communities: 22<br>Modularity: 0.7483 |
| 51 | Earth Microbiome Project | LC-MS | - | MSV000083475 | https://gnps.ucsd.edu/ProteoSAFe/status.jsp?task=9ec6a7944b7441b385b8c3080e6c1174 https://gnps.ucsd.edu/ProteoSAFe/status.jsp?task=e7de3dcb30e54d8eaec2e6070b2e7d6d | Entire graph: 16182 nodes<br>GCC: 14940 nodes<br>Density: 0.0023<br>Communities: 23<br>Modularity: 0.7530 |
| 52 | Bile Acid Gavaged Mice | LC-MS | - | MSV000083004 | https://gnps.ucsd.edu/ProteoSAFe/status.jsp?task=eefbf60290d843028425d1bf4f1de618 https://gnps.ucsd.edu/ProteoSAFe/status.jsp?task=f4826b7ce0b745f2a768c5c8338cb1cd | Entire graph: 10415 nodes<br>GCC: 9707 nodes<br>Density: 0.0034<br>Communities: 31<br>Modularity: 0.8385 |
| 53 | T. Septentrionalis Garden Extracts | LC-MS | - | MSV000084024 | https://gnps.ucsd.edu/ProteoSAFe/status.jsp?task=e649468df3624de89c2073747a5b02f0 https://gnps.ucsd.edu/ProteoSAFe/status.jsp?task=e2bc5bee49b04a1aa3eb785d4aad32d5 | Entire graph: 6708 nodes<br>GCC: 5904 nodes<br>Density: 0.0041<br>Communities: 32<br>Modularity: 0.8260 |
| 54 | Human Fecal Samples | LC-MS | - | MSV000091381 | https://gnps.ucsd.edu/ProteoSAFe/status.jsp?task=81a485053c1b4c44a5aff14d5f88837e https://gnps.ucsd.edu/ProteoSAFe/status.jsp?task=dd12df9affd14dc09a82c57959b79efa | Entire graph: 10848 nodes<br>GCC: 10263 nodes<br>Density: 0.0033<br>Communities: 30<br>Modularity: 0.7318 |
| 55 | Skin Swabs for Microbial Metabolism Study | Positive Mode LC-MS (C18) | - | MSV000088283 | https://gnps.ucsd.edu/ProteoSAFe/status.jsp?task=d9526ac893a64553830d53d98e49b498 https://gnps.ucsd.edu/ProteoSAFe/status.jsp?task=4684f5609c79471fbaa126ef237fe907 | Entire graph: 7558 nodes<br>GCC: 7353 nodes<br>Density: 0.0066<br>Communities: 20<br>Modularity: 0.7075 |
| 56 | Skin Swabs for Microbial Metabolism Study | Negative Mode LC-MS (C18) | - | MSV000088283 | https://gnps.ucsd.edu/ProteoSAFe/status.jsp?task=c886d37d6e3a4324ad306c13c54d4acb https://gnps.ucsd.edu/ProteoSAFe/status.jsp?task=de44d73ff872478c9c8e36eb12ccd835 | Entire graph: 2150 nodes<br>GCC: 1943 nodes<br>Density: 0.0129<br>Communities: 12<br>Modularity: 0.6310 |
| 57 | Skin Swabs for Microbial Metabolism Study | Positive Mode LC-MS (HILIC) | - | MSV000088283 | https://gnps.ucsd.edu/ProteoSAFe/status.jsp?task=5f99cbdaa69b4d68b520c372f89cd2e7 https://gnps.ucsd.edu/ProteoSAFe/status.jsp?task=6181f073bcfd4a57866dfbf9950a3592 | Entire graph: 3344 nodes<br>GCC: 3162 nodes<br>Density: 0.0093<br>Communities: 14<br>Modularity: 0.7485 |
| 58 | Skin Swabs for Microbial Metabolism Study | Negative Mode LC-MS (HILIC) | - | MSV000088283 | https://gnps.ucsd.edu/ProteoSAFe/status.jsp?task=50258745f69946288aa63af421af3806 https://gnps.ucsd.edu/ProteoSAFe/status.jsp?task=517b145c259c4f16af944d043e8f2781 | Entire graph: 2742 nodes<br>GCC: 2543 nodes<br>Density: 0.0078<br>Communities: 15<br>Modularity: 0.7152 |
| 59 | Tomato Endophyte | LC-MS | - | MSV000081463 | https://gnps.ucsd.edu/ProteoSAFe/status.jsp?task=9544958157d247f3abb8da64a7e78bb1 https://gnps.ucsd.edu/ProteoSAFe/status.jsp?task=2ad66e3e4b0b4bc7a587c4daad2131e6 | Entire graph: 11105 nodes<br>GCC: 10607 nodes<br>Density: 0.0033<br>Communities: 21<br>Modularity: 0.7007 |
| 60 | Lichen Isolates | LC-MS | - | MSV000080115 | https://gnps.ucsd.edu/ProteoSAFe/status.jsp?task=7e83b932c79e423cb35eea3c96fe94c6 https://gnps.ucsd.edu/ProteoSAFe/status.jsp?task=a84f71928ecf4b43aa6c7522324d9770 | Entire graph: 1799 nodes<br>GCC: 618 nodes<br>Density: 0.0398<br>Communities: 11<br>Modularity: 0.4617 |
| 61 | NIH Pharmacologically Active Library | LC-MS | - | MSV000078710 | https://gnps.ucsd.edu/ProteoSAFe/status.jsp?task=4ddf2c70412744b89a9a96209d0ac975 https://gnps.ucsd.edu/ProteoSAFe/status.jsp?task=1f43d8ef135a498fa800eb5a454cec89 | Entire graph: 2391 nodes<br>GCC: 1046 nodes<br>Density: 0.0282<br>Communities: 18<br>Modularity: 0.6977 |
| 62 | NIH Natural Products Library | LC-MS | - | MSV000078708 | https://gnps.ucsd.edu/ProteoSAFe/status.jsp?task=f89ee722a41f4c19ac5117a136f4372d https://gnps.ucsd.edu/ProteoSAFe/status.jsp?task=eb8e895325b540809a9b9ca9c651dbc6 | Entire graph: 2489 nodes<br>GCC: 1573 nodes<br>Density: 0.0084<br>Communities: 27<br>Modularity: 0.8203 |
| 63 | Milk and Yogurt Metabolite Analysis | LC-MS | - | MSV000082387 | https://gnps.ucsd.edu/ProteoSAFe/status.jsp?task=c97bbabf48d248cc9695c46975a263fc https://gnps.ucsd.edu/ProteoSAFe/status.jsp?task=8ddde51fe41844619426fc36b24e3255 | Entire graph: 3069 nodes<br>GCC: 2817 nodes<br>Density: 0.0091<br>Communities: 23<br>Modularity: 0.8292 |
| 64 | Metabolite Analysis of Meat Samples | LC-MS | - | MSV000082423 | https://gnps.ucsd.edu/ProteoSAFe/status.jsp?task=dc02e34956d74cffabf5786e79453f50 https://gnps.ucsd.edu/ProteoSAFe/status.jsp?task=d50290f2a3da47648f1a10dae998be8b | Entire graph: 2770 nodes<br>GCC: 2500 nodes<br>Density: 0.0109<br>Communities: 24<br>Modularity: 0.6428 |
| 65 | Metabolite Analysis of Tea Samples | LC-MS | - | MSV000082388 | https://gnps.ucsd.edu/ProteoSAFe/status.jsp?task=4b27a3df62c0441c9dce2c53fdc6c877 https://gnps.ucsd.edu/ProteoSAFe/status.jsp?task=f71f15f5be0e4a19a05ce35dd105a3ed | Entire graph: 1200 nodes<br>GCC: 1018 nodes<br>Density: 0.0214<br>Communities: 17<br>Modularity: 0.7281 |
| 66 | Bile Acid Modification of Mouse and Carnivore Fecal Samples | LC-MS | - | MSV000091663 | https://gnps.ucsd.edu/ProteoSAFe/status.jsp?task=3581c7e9c5ef4eb1a92f5981c7d90c81 https://gnps.ucsd.edu/ProteoSAFe/status.jsp?task=80ead08fa2a1496999c9943bc6241d7d | Entire graph: 5266 nodes<br>GCC: 4389 nodes<br>Density: 0.0083<br>Communities: 24<br>Modularity: 0.8163 |
| 67 | Metabolite Analysis of Coffee Samples | LC-MS | - | MSV000082386 | https://gnps.ucsd.edu/ProteoSAFe/status.jsp?task=bbb12c5856fb4a57a1bc2c3959030474 https://gnps.ucsd.edu/ProteoSAFe/status.jsp?task=cbe3f2b5f7c04c809a7241f4ff458f6f | Entire graph: 935 nodes<br>GCC: 645 nodes<br>Density: 0.0195<br>Communities: 21<br>Modularity: 0.6470 |
| 68 | Coral Extracts | LC-MS | - | MSV000081908 | https://gnps.ucsd.edu/ProteoSAFe/status.jsp?task=074dbb782bca4bc79f4e0bdb8686a054 https://gnps.ucsd.edu/ProteoSAFe/status.jsp?task=ec5c914ae7344cf8b7a1b9a798a38b59 | Entire graph: 2122 nodes<br>GCC: 1694 nodes<br>Density: 0.0119<br>Communities: 18<br>Modularity: 0.7837 |
| 69 | Metabolite Analysis of Tomato Samples | LC-MS | - | MSV000082391 | https://gnps.ucsd.edu/ProteoSAFe/status.jsp?task=30821b0c6b094010b3b42efcb425a604 https://gnps.ucsd.edu/ProteoSAFe/status.jsp?task=8ffc0692adb54101896923d38f761755 | Entire graph: 1571 nodes<br>GCC: 1278 nodes<br>Density: 0.0156<br>Communities: 20<br>Modularity: 0.7518 |
| 70 | Metabolic Profiles of Water Samples | LC-MS | - | MSV000080015 | https://gnps.ucsd.edu/ProteoSAFe/status.jsp?task=19d59a42e9aa4ce39e11fcad5d93f98c https://gnps.ucsd.edu/ProteoSAFe/status.jsp?task=2b6993f19e604e3692d12df2ac49839e | Entire graph: 5116 nodes<br>GCC: 4747 nodes<br>Density: 0.0075<br>Communities: 18<br>Modularity: 0.7521 |
| 71 | Cultures of Paenibacillus alevei in Spent Coffee Grounds | LC-MS | - | MSV000087141 | https://gnps.ucsd.edu/ProteoSAFe/status.jsp?task=5bfc1afb85584a05883e1605d2539f7d https://gnps.ucsd.edu/ProteoSAFe/status.jsp?task=31117bdab0ee4e9fb7b0e273f1294e13 | Entire graph: 1237 nodes<br>GCC: 813 nodes<br>Density: 0.0332<br>Communities: 10<br>Modularity: 0.4980 |
