## Supplemental Table 2 for "Ordering molecular diversity in untargeted metabolomics via molecular community networking"

| Massive ID | DOI | Massive Link |
| --- | --- | --- |
| MSV000084475 | doi:10.25345/C5JH3S | https://massive.ucsd.edu/ProteoSAFe/dataset.jsp?task=295cb2aeccef486583a2e36f14ff9713 |
| MSV000081351 | N/A | https://massive.ucsd.edu/ProteoSAFe/dataset.jsp?task=8caf4ebc01bf4cbdb04fbb1b5922ec8e |
| MSV000083024 | N/A | https://massive.ucsd.edu/ProteoSAFe/dataset.jsp?task=d79ac6a8331940ee8d3bb9f7715a7b55 |
| MSV000084218 | doi:10.25345/C53W9T | https://massive.ucsd.edu/ProteoSAFe/dataset.jsp?task=be1f8c47243b492686df4423dab8f50e |
| MSV000084314 | doi:10.25345/C5WQ0T | https://massive.ucsd.edu/ProteoSAFe/dataset.jsp?task=25cc4f9135c6428aabe1f41a9e54c369 |
| MSV000079598 | N/A | https://massive.ucsd.edu/ProteoSAFe/dataset.jsp?task=856f31ce9d6c41e1a410fd127508959b |
| MSV000080469 | N/A | https://massive.ucsd.edu/ProteoSAFe/dataset.jsp?task=51f7681681c9433db2671f8b028c0b93 |
